# Structure of a TRAPPII-Rab11 activation intermediate reveals GTPase substrate selection mechanisms

**DOI:** 10.1101/2021.12.20.473486

**Authors:** Saket R. Bagde, J. Christopher Fromme

**Affiliations:** Department of Molecular Biology and Genetics, Weill Institute for Cell and Molecular Biology, Cornell University, Ithaca, NY 14853 USA

## Abstract

Rab1 and Rab11 are essential regulators of the eukaryotic secretory and endocytic recycling pathways. The TRAPP complexes activate these GTPases via nucleotide exchange using a shared set of core subunits. The basal specificity of the TRAPP core is towards Rab1, yet the TRAPPII complex is specific for Rab11. A steric gating mechanism has been proposed to explain TRAPPII counterselection against Rab1. Here we present cryoEM structures of the 22-subunit TRAPPII complex from budding yeast, including a TRAPPII-Rab11 nucleotide exchange intermediate. The Trs130 subunit provides a “leg” that positions the active site distal to the membrane surface, and this leg is required for steric gating. The related TRAPPIII complex is unable to activate Rab11 due to a repulsive interaction, which TRAPPII surmounts using the Trs120 subunit as a “lid” to enclose the active site. TRAPPII also adopts an open conformation enabling Rab11 to access and exit from the active site chamber.

## Introduction

Protein and membrane traffic in eukaryotic cells is controlled by Rab GTPases that function by recruiting effector protein machinery to generate, transport, and tether membrane vesicles and tubules (*1, 2*). Rab1 and Rab11 have been described as the gatekeepers of the Golgi complex (*3*). Rab1 and its close paralogs enable traffic to enter the Golgi from the endoplasmic reticulum by recruiting vesicle tethers (*4*– *6*). Rab11 and its close paralogs regulate anterograde traffic out of Golgi and recycling endosome compartments by recruiting effectors to make and transport vesicles (*7*–*10*). Targeted activation of Rab1 and Rab11 at distinct compartments is therefore an essential feature of eukaryotic cells.

The master regulators of Rab GTPase pathways are the GEFs (<u>g</u>uanine-nucleotide <u>e</u>xchange factor) that must decide where and when to activate their substrate GTPases. There are at least three reported GEFs for Rab11 (*3, 11*–*15*); the Rab11 GEF distributed most broadly throughout the eukaryotic kingdom appears to be the multi-subunit TRAPPII complex (*16, 17*). TRAPPII shares a set of core subunits with the related TRAPPIII complex, yet the TRAPPIII complex activates a different substrate, Rab1 (*18, 19*). Mutations in TRAPP subunits are known to be associated with a diverse array of human diseases (*20*).

Previous work using the budding yeast (*S. cerevisiae*, hereafter “yeast”) model determined that the same active site was used by both TRAPP complexes (*21, 22*), and that the C-terminal HVD (hypervariable domain) tails of the two Rabs play a key role in TRAPP complex specificity (*23*). As the basal specificity of the TRAPP core is towards Rab1, it was hypothesized that TRAPPII-specific subunits must make additional contact with Rab11 to enable its activation. In contrast, TRAPPII counterselection against Rab1 was proposed to be enforced via a steric gating mechanism, in which the shorter Rab1 HVD tail prevented Rab1 from accessing the TRAPPII active site. The key finding supporting the steric gating model was the observation that addition of a ∼10 residue Gly-Ser linker to the Rab1 HVD tail enabled Rab1 to be activated by TRAPPII both *in vitro* and *in vivo (23)*.

Earlier studies led to an atomic model of the core TRAPP complex and revealed the structural basis for Rab1 nucleotide exchange (*22, 24*). Recently, cryoEM structures of the intact yeast and fly TRAPPIII complexes were reported (*25, 26*); these studies elucidated the architecture of TRAPPIII and determined how it binds membranes during the Rab activation reaction. Low resolution structures of TRAPPII from multiple organisms have also been determined (*26*–*29*), yet the central question of how the same active site can activate different Rab substrates in different contexts remains unresolved. It is not known why TRAPPIII does not activate Rab11, how TRAPPII is able to activate Rab11, or how TRAPPII counterselects against Rab1 by steric gating.

Here we report cryoEM structures of the yeast TRAPPII complex, including the structure of a TRAPPII-Rab11 activation intermediate. These structures reveal the architecture of the complex, how the complex binds and activates Rab11, and the orientation of the complex on the membrane. We use the structure to guide functional experiments that further determine why TRAPPIII is unable to activate Rab11, and why TRAPPII is unable to activate Rab1. We describe two different conformations of TRAPPII: an open state that allows Rab11 to access the active site chamber, and a closed state in which the Trs120 lid encloses the chamber to enable Rab11 activation. Our findings provide a structural and mechanistic understanding of how distinct GTPases can be differentiated by different protein complexes sharing a common active site.

## Results

### Molecular architecture of the yeast TRAPPII complex

Yeast TRAPPII comprises 22 subunits encoded by 10 genes, and yeast have two Rab11 paralogs, named Ypt31 and Ypt32. We purified endogenous TRAPPII from yeast and prepared a stable TRAPPII-Rab11 complex by incubating purified TRAPPII with purified Rab11/Ypt32 in the presence of alkaline phosphatase (fig. S1). We then used single-particle cryoEM to determine structures of TRAPPII by itself and bound to Rab11/Ypt32 (figs. S2, S3 and S4 and Table S1). Due to flexibility of the complex and the presence of multiple conformational states, we made extensive use of symmetry expansion, focused refinement, and 3D classification approaches. The resulting focused reconstructions had 0.143-FSC resolutions ranging from 3.4Å to 3.9Å. The resolutions of the consensus monomer and dimer reconstructions were 3.7Å and 4.1Å, respectively. To facilitate interpretation and model building of the entire complex, we produced composite maps using density-modified reconstructions of the focused refinements. We were able to confidently build over 84% of the residues in the complex, representing virtually all of the conserved elements and domains.

Within the dimeric TRAPPII particles, individual monomers exhibited two major conformations which we refer to as the open and closed states. TRAPPII monomers bound to Rab11 adopted only the closed conformation. We initially focus our description and analysis on the Rab11-bound closed conformation and will describe the open conformation further below.

Viewed from the “top”, the shape of the TRAPPII dimer is reminiscent of a butterfly (Fig. 1, A and B). Each of the 11-subunit monomers forms a triangle shape in which the TRAPP core subunits appear to be grasped by tongs composed of the TRAPPII-specific Trs120 and Trs130 subunits. The arrangement and structures of the core subunits are essentially identical to their arrangement and structure within the TRAPPIII and isolated TRAPP core complexes (*22, 24*–*26*). In the TRAPPII structure Trs130 is linked to the core via Tca17, which serves as a bridge to the Trs33 core subunit, consistent with previous biochemical analyses (*30, 31*). Trs120 connects to the core by binding to Trs20 at the site of a human disease mutation that is known to disrupt both TRAPPII and TRAPPIII assembly (*32*–*34*). The arrangement of Trs120 and Trs130 in the complex relative to the core (figs. S5 and S6) fits well with published experimental data (*30*– *34*), but is different from that inferred from negative stain analysis reported previously (*28*). Trs65 forms an extensive interface between the two monomers (fig. S7), confirming its established role in dimerization of the TRAPPII complex (*28, 30*). As the structures of Trs65, Trs120, and Trs130 have not been previously reported, we provide more details regarding their folds and comparisons to structural homologs in figs. S5, S6, and S7.

**Fig. 1.**
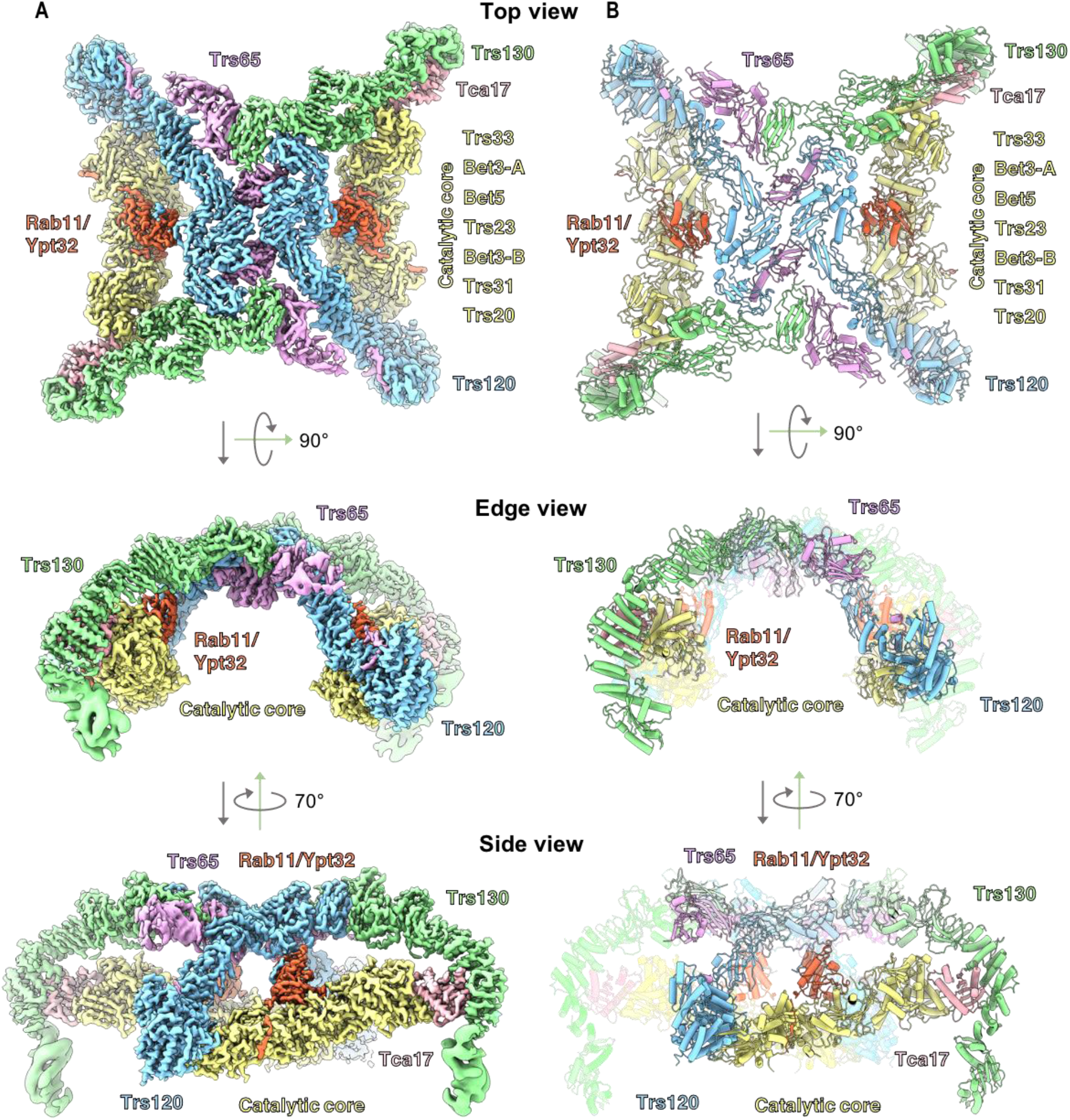
CryoEM structure of a TRAPPII-Rab11/Ypt32 activation intermediate. (**A**) cryoEM map of the TRAPPII-Rab11/Ypt32 complex. Subunits are labeled and the complex is shown from “top”, “edge”, and “side” views. (**B**) Atomic model of the complex.

Viewed from the “edge”, TRAPPII is shaped like an arch. This curved shape was not observed in published negative stain analysis, perhaps due to staining artefacts(*28*). Viewed from the “side”, each monomer forms a chamber in which Rab11 is sandwiched between the core and Trs120. The N-terminal portion of Trs120 contains a GTPase fold and bears a strong resemblance to the TRAPPIII subunit Trs85. Trs120 interacts with Trs20 in a manner that is remarkably similar to the interaction between Trs85 and Trs20 in TRAPPIII (fig. S5).

Trs130 interacts with Tca17 (fig. S6) in a manner identical to that recently predicted computationally (*35*), at the site of a mutation in the human Tca17 paralog TRAPPC2L that disrupts this interaction and is associated with a neurodevelopmental disorder (*36*). Strikingly, the N-terminal region of Trs130 extends a significant distance down below the core of the complex, and we refer to this extension as a “leg”. This leg region appeared to be somewhat flexible, requiring our use of focused refinements to produce reasonable density maps that facilitated model building. The “foot” of this leg adopts a GTPase fold, akin to the GTPase folds also found in Trs120 of TRAPPII and Trs85 of TRAPPIII, although we note that none of these GTPase-like domains appear able to bind nucleotide. We describe the importance of the Trs130 leg region further below.

### Interaction with Rab11 and orientation of TRAPPII on the membrane

As expected from previous work, the nucleotide-binding domain (NBD) of Rab11/Ypt32 binds to the TRAPP core in the same active site used to bind the Rab1/Ypt1 NBD (*21, 22*) (Fig. 2 and, fig. S8A). GEFs exchange nucleotide by destabilizing nucleotide binding to the GTPase, resulting in a nucleotide-free, GEF-bound intermediate. Superposition of the published structure of inactive, GDP-bound Rab11/Ypt32 (*37*) onto the nucleotide-free activation intermediate indicates that the GTPase “switch I” region has undergone a dramatic opening concomitant with nucleotide release (Fig. 2A). To compare TRAPP activation of Rab1 versus Rab11, we superimposed TRAPPII-bound Rab11/Ypt32 onto the published structures of Rab1/Ypt1 bound to the isolated TRAPP core (*22*) (fig. S8A) and Rab1/Ypt1 bound to TRAPPIII (*25*) (fig. S8B). This comparison indicates the structures of the Rab1 and Rab11 nucleotide-free intermediates, and their interfaces with the TRAPP core of their corresponding GEF, are quite similar. As found for Rab1/Ypt1, Rab11/Ypt32 makes contact with the Bet3, Bet5, Trs31 and Trs23 subunits, and we observed density corresponding to the C-terminus of one of the Bet3 subunits interacting with the opened nucleotide binding site. Taken together, these observations suggest that the core active sites of both TRAPPII and TRAPPIII use the same mechanism to disrupt nucleotide binding as was first determined for the isolated TRAPP core (*22*).

**Fig. 2.**
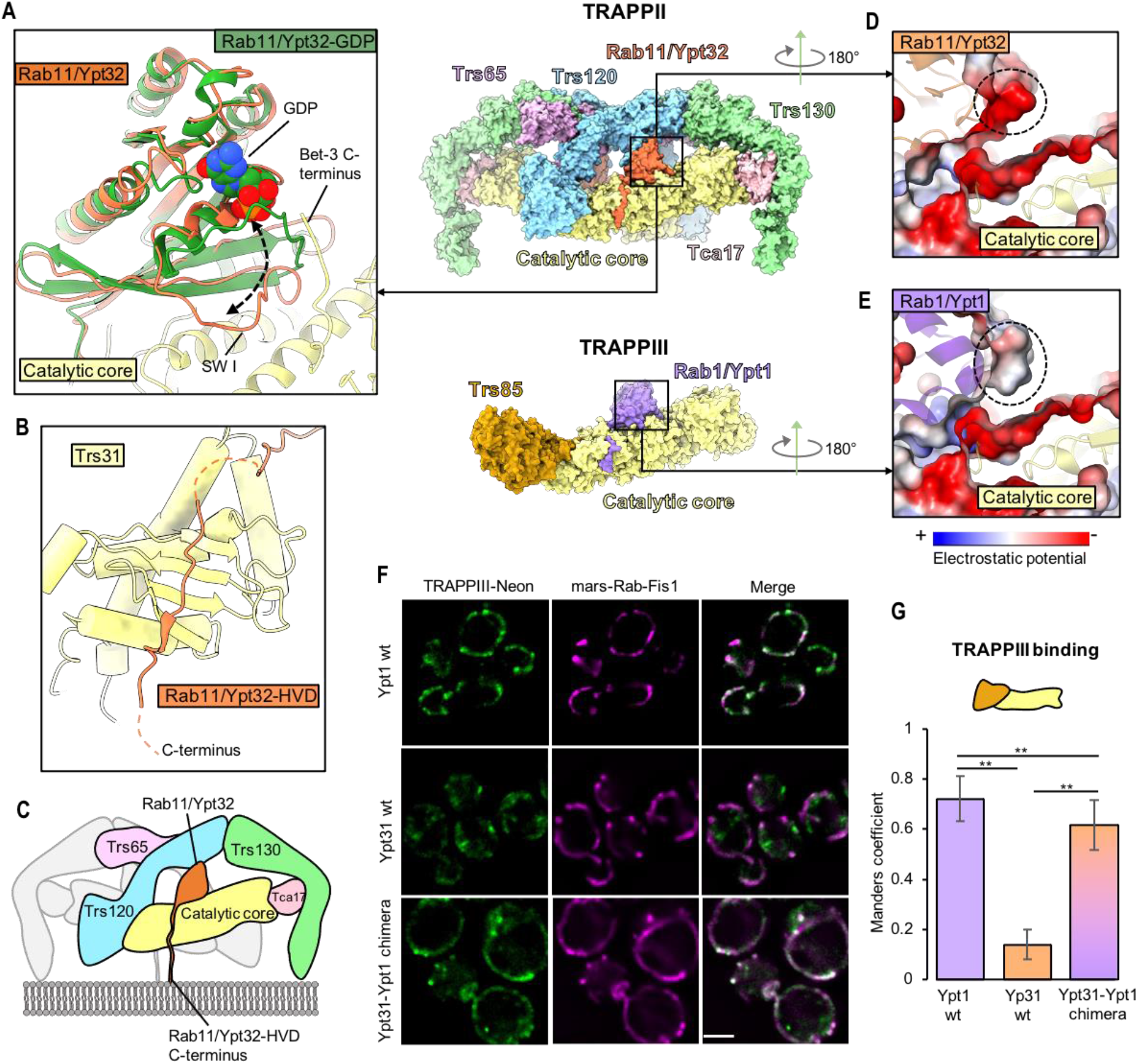
Interactions between the TRAPPII core and Rab11/Ypt32. (**A**) Close-up view of the interaction between the catalytic core (yellow) and nucleotide-free Rab11/Ypt32 (orange). The structure of inactive, GDP-bound Rab11/Ypt32(*37*) is superimposed to show the conformational changes associated with nucleotide release. (**B**) Close-up view of the Rab11/Ypt32 HVD binding site on the Trs31 core subunit. (**C**) Model for orientation of the complex on the membrane surface. (**D**) Close-up view of TRAPPII-bound Rab11/Ypt32, colored by electrostatics. The anionic patch on the Rab is highlighted with a dashed circle. (**E**) Close-up view of TRAPPIII-bound Rab1/Ypt1(*25*), colored by electrostatics. The equivalent patch, which is more neutral in Rab1/Ypt1, is highlighted with a dashed circle. (**F**) Imaging data from a GRab-IT experiment testing association of indicated Rab constructs with TRAPPIII. Note all Rab constructs harbor a mutation that prevents nucleotide binding (see methods). Scale bar shown is 2µm. (**G**) Quantification of the GRab-IT data.

Somewhat unexpectedly, we observed that the HVD of Rab11/Ypt32 binds to the same pocket on the Trs31 core subunit (Fig. 2B) that binds to the HVD of Rab1/Ypt1 in TRAPPIII (25). Given the two-fold symmetry of the TRAPPII complex and the fact that activated Rab11 is anchored to the membrane via hydrophobic prenyl modifications of C-terminal cysteine residues, we can confidently predict the orientation of the complex on the membrane surface (Fig. 2C).

### Rab11 is a poor substrate for the TRAPP core due to an unfavorable interaction

To understand why Rab11 is a poor substrate of the TRAPP core, we compared the core-Rab1 and core-Rab11 interaction interfaces. We identified a potentially unfavorable repulsive interaction of the core with Rab11 due to a negatively charged surface of Rab11 (Fig. 2D and fig. S8, C and D). The corresponding surface of Rab1 is less anionic and therefore expected to be more favorable (Fig. 2E and fig. S8, E and F). To test whether this surface of Rab11 was responsible for the inability of the TRAPP core to activate Rab11, we grafted the corresponding surface of Rab1 onto Rab11. We found that this grafted Rab chimera (fig. S8G) gained the ability to interact stably with TRAPPIII in the “GRab-IT” (<u>G</u>EF-<u>Rab</u> Interaction <u>T</u>est) ectopic localization assay (*23*), in which Rab11 does not normally interact with TRAPPIII (Fig. 2, F and G). This indicates that Rab11 is a poor substrate of the core due, at least in part, to this unfavorable interaction surface.

### Trs120 forms a lid that encloses the active site chamber

Our group previously proposed that TRAPPII is able to activate Rab11 because the TRAPPII-specific subunits provide additional unknown interaction(s) with Rab11 (*21, 23*). The structure of the TRAPPII-Rab11/Ypt32 activation intermediate supports this proposal, as we now observe that Trs120 makes multiple direct contacts with Rab11/Ypt32 on a surface of the GTPase that is distal to its interaction with the core (Fig. 3A). We identified three different ordered loops of Trs120 that contact Rab11. One of these loops (“loop 1”) interacts with Rab11/Ypt32 by adopting a β-strand that binds to the edge of the GTPase β-sheet (Fig. 3A). The other loops (“loop 2” and “loop 3”) make primarily electrostatic interactions with Rab11/Ypt32 (Fig. 3B).

**Fig. 3.**
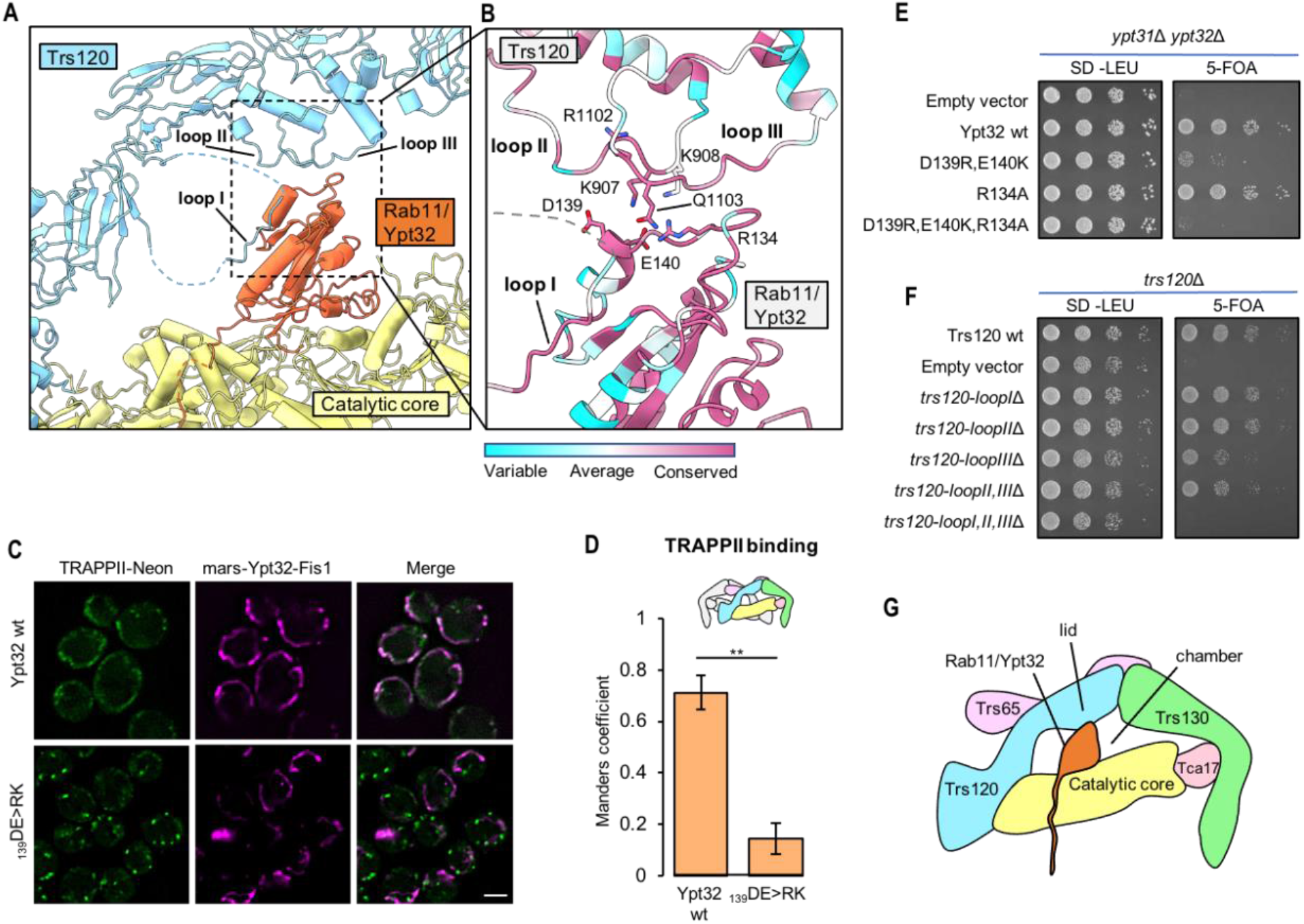
Trs120 forms a lid required for Rab11/Ypt32 interaction. (**A**) View of the Rab11/Ypt32 binding interface with Trs120. Ordered loops that contact Rab11/Ypt32 are indicated. (**B**) Close-up view of (A), with residue conservation indicated. (**C**) Imaging data from a GRab-IT experiment monitoring the association between TRAPPII and nucleotide-free wild-type or mutant Rab11/Ypt32. Note both “wild-type” and mutant Rab11/Ypt32 constructs harbor a mutation that prevents nucleotide binding (see methods). (**D**) Quantitation of the data in (C). (**E**) Complementation test assessing the effects of residue substitutions in Rab11/Ypt32 on cell viability. (**F**) Complementation test assessing the effects of loop-deletion mutants of Trs120 on cell viability. (**G**) Cartoon schematic illustrating the interaction of Rab11/Ypt32 with the TRAPPII monomer. Trs120 forms a lid to enclose Ypt32 within the active site chamber.

To determine the importance of the interactions between Trs120 and Rab11, we tested mutants in conserved residues that we expected might disrupt this interaction. In the GRab-IT assay, charge-reversal mutants of Rab11/Ypt32 expected to introduce repulsive interactions lost the ability to interact stably with Trs120 (Fig. 3, C and D). These mutants were also unable to provide the essential function of the *YPT31/32* genes in a complementation assay (Fig. 3E and fig. S10A), although we cannot rule out the possibility that their loss of function was due to disruption of effector binding. We therefore tested the ability of *trs120* loop-deletion mutants to complement the essential function of the *TRS120* gene. We found that deletion of any single loop resulted in only a minor growth phenotype, while deletion of all three loops resulted in a significant growth defect (Fig. 3F and fig. S10, B and C). Our interpretation of this set of results is that, although some specificity is likely contributed by these interactions, these ordered loops may be more important for providing steric bulk than for providing specific interactions with Rab11. We therefore propose that Trs120 serves as a lid to enclose the active site, creating an active site chamber (Fig. 3G). By lowering the off-rate of Rab11 from the TRAPP core, the Trs120 lid may enable TRAPPII to catalyze nucleotide exchange in spite of the otherwise unfavorable interaction between Rab11 and the core described above.

### Trs130 provides a leg that lifts the active site above the membrane to enforce steric gating

The cryoEM density for the portion of the Rab11/Ypt32 HVD bound to the TRAPP core was clear enough that we could confidently model HVD residues 202-207 bound to Trs31 (fig. S8H), and this modeled sequence is consistent with the established HVD sequence requirements for Rab11 activation by TRAPPII (*23*). The residues of the Rab1/Ypt1 HVD required for binding to the equivalent pocket of Trs31 in TRAPPIII have also been mapped (*25*). There is little sequence homology shared between the portions of the Rab1 and Rab11 HVDs bound to Trs31 (fig. S8I), yet both sequences include the “CIM” motif required for post-translational prenylation (*38*). Furthermore, in both cases the HVD adopts a β-strand structure that makes backbone contacts with the β-sheet in Trs31 (fig. S8H and reference (*25*)).

The relative positions of the HVD residues bound to Trs31 in each Rab-TRAPP complex (fig. S8I) supports the steric gating model in which the Rab1 HVD is not long enough for the Rab1 NBD to reach the TRAPPII active site (Fig. 4A). To continue probing the steric gating model, we aimed to further test the HVD length requirement of Rab11. Previous analysis of Rab11 tested the impact of truncations of a more N-terminal portion of the HVD, residues 186-200 (reference (*23*)), that our structure reveals to lie in between the Trs31-binding sequence (residues 202-207) and the NBD. We therefore produced a new truncation of the HVD, removing residues 208-215, which are C-terminal to the Trs31-binding sequence and are therefore expected to be required for spanning the gap between Trs31 and the membrane surface. This 8-residue truncation resulted in a loss of cell viability, which was rescued by substitution of the truncated residues with an 8-residue Gly-Ser linker (Fig. 4B). The truncation lost its ability to interact with TRAPPII in the GRab-IT assay, but the interaction was restored by the same Gly-Ser linker (Fig. 4, C and D). The 8-residue truncation also prevented activation of fluorescently-tagged Rab11/Ypt32 in cells, as monitored by its colocalization with the late-Golgi marker Sec7, and this phenotype was also rescued by addition of an 8-residue Gly-Ser linker (fig. S10D). These data provide additional evidence supporting the existence of an HVD length constraint for Rab activation by TRAPPII.

**Fig. 4.**
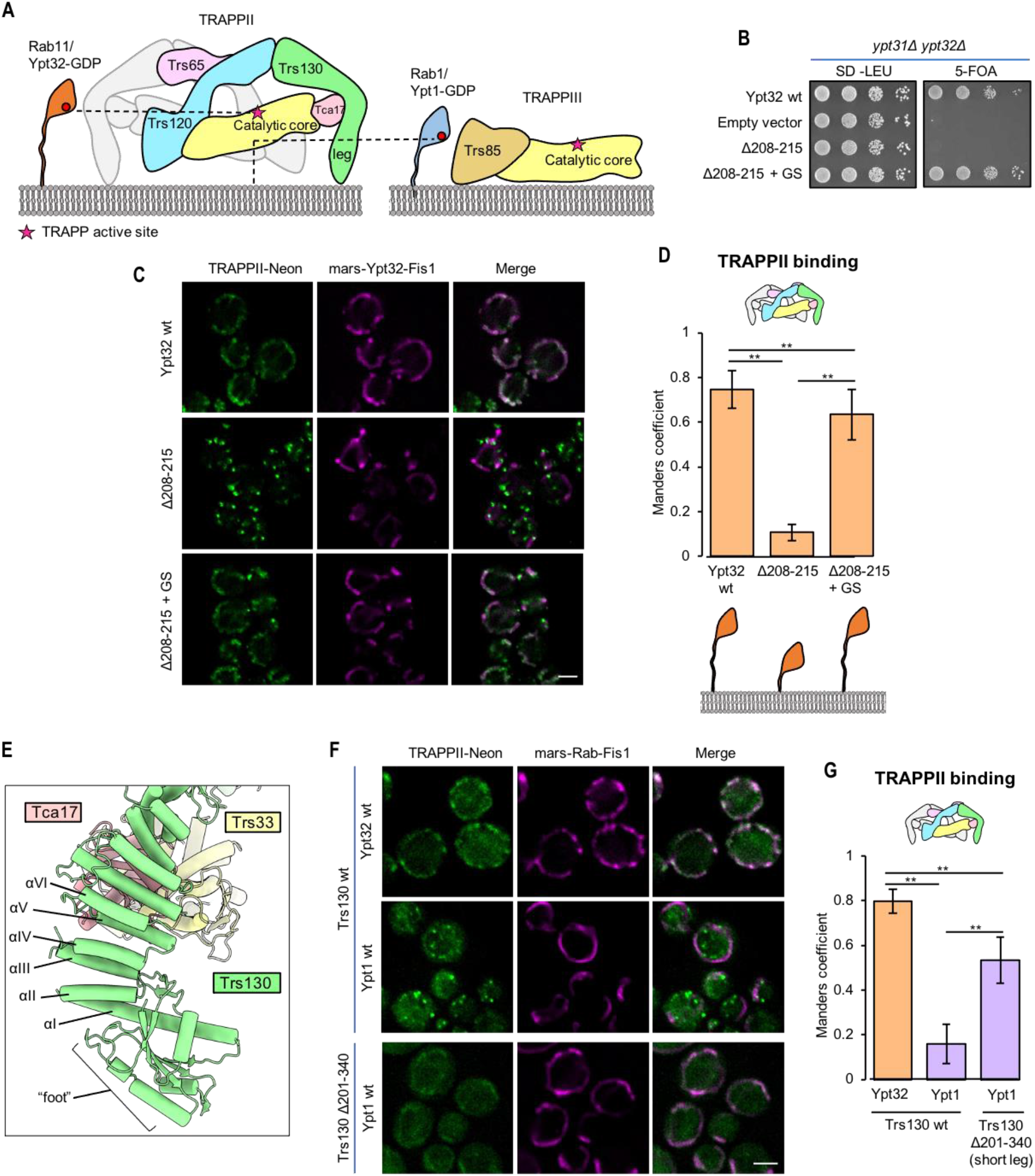
The Trs130 “leg” enforces counterselection against Rab1 by steric gating. (**A**) Schematic illustrating the steric gating mechanism, in which the Rab1 HVD is not long enough to enable access to the TRAPPII active site. (**B**) Complementation test assessing the effects of Rab11/Ypt32 HVD truncation and residue substitution on cell viability. (**C**) GRab-IT imaging data to test the interaction of TRAPPII with the same Rab11/Ypt32 constructs tested in (B). (**D**) Quantitation of the data in (C). (**E**) View of the Trs130 leg structure. (**F**) GRab-IT imaging data to test the interaction of Rab11/Ypt32 with TRAPPII harboring the “short-leg” Trs130 construct. (**G**) Quantitation of the data in (F). Scale bars shown in (C) and (F) are 2µm.

The N-terminal “leg” of Trs130 appeared likely to serve as a structural element responsible for lifting the active site away from the membrane (Fig. 4A). To test this possible role of the Trs130 leg, we designed an internal truncation of this leg that preserved the putative membrane-binding “foot” region, which includes the GTPase-like domain, at the N-terminus of Trs130 (Fig. 4E). The truncation removed 5 *α*-helices (residues 201-340, helices I-IV) from the *α*-solenoid portion of Trs130 to generate a “short-leg” construct. As predicted by the steric gating hypothesis, truncation of the Trs130 leg resulted in a gain-of-function phenotype: the short-leg mutant gained the ability to stably interact with Rab1/Ypt1 in the GRab-IT assay (Fig. 4, F and G). Therefore, the Trs130 leg is required to enforce counter-selection against Rab1.

### TRAPPII conformational change may enable Rab11 to access the active site

A mixture of open and closed states were present in the TRAPPII-only (lacking Rab11/Ypt32) cryoEM data, and 3D classification resulted in classes representing four different conformations of the TRAPPII dimer (fig. S3): ∼30% of particles sorted into a class in which both monomers were closed (“closed/closed”); ∼50% of particles sorted into two classes in which one monomer was closed and the other monomer was either open (“closed/open”) or partially open (“closed/partially-open”, a state with a conformation that is intermediate between the open and closed states); ∼20% of particles sorted into a class in which one monomer was open and the other monomer was partially open (“open/partially-open”). A state resembling the closed/open state appears to have also been previously observed in a subset of particles imaged during negative-stain EM analysis of TRAPPII (*28*).

Classification of the TRAPPII-Rab11/Ypt32 cryoEM data resulted in three different classes (fig. S2): ∼30% of particles sorted into the closed/closed state, ∼40% of particles sorted into the closed/partially-open state, and ∼30% of particles sorted into the closed/open state. Importantly, Rab11/Ypt32 was not bound to the open or partially-open monomers in any of the classes. This is consistent with the idea that closing of the Trs120 lid is required for productive binding and subsequent nucleotide exchange of Rab11.

Further 3D classification of the closed/open class of the TRAPPII-Rab11/Ypt32 cryoEM data resulted in a subset (∼50%) of the particles sorting into a class in which Rab11/Ypt32 bound at full occupancy to the closed monomer (fig. S2). Among classes containing at least one open or partially-open monomer, this class resulted in the highest resolution reconstruction (4.5 Å). We therefore used this class for further focused refinements and obtained focused maps ranging in overall resolution values of 3.8-4.2Å which aided in model building and analysis of the open conformational state (Fig. 5, A and B).

**Fig. 5.**
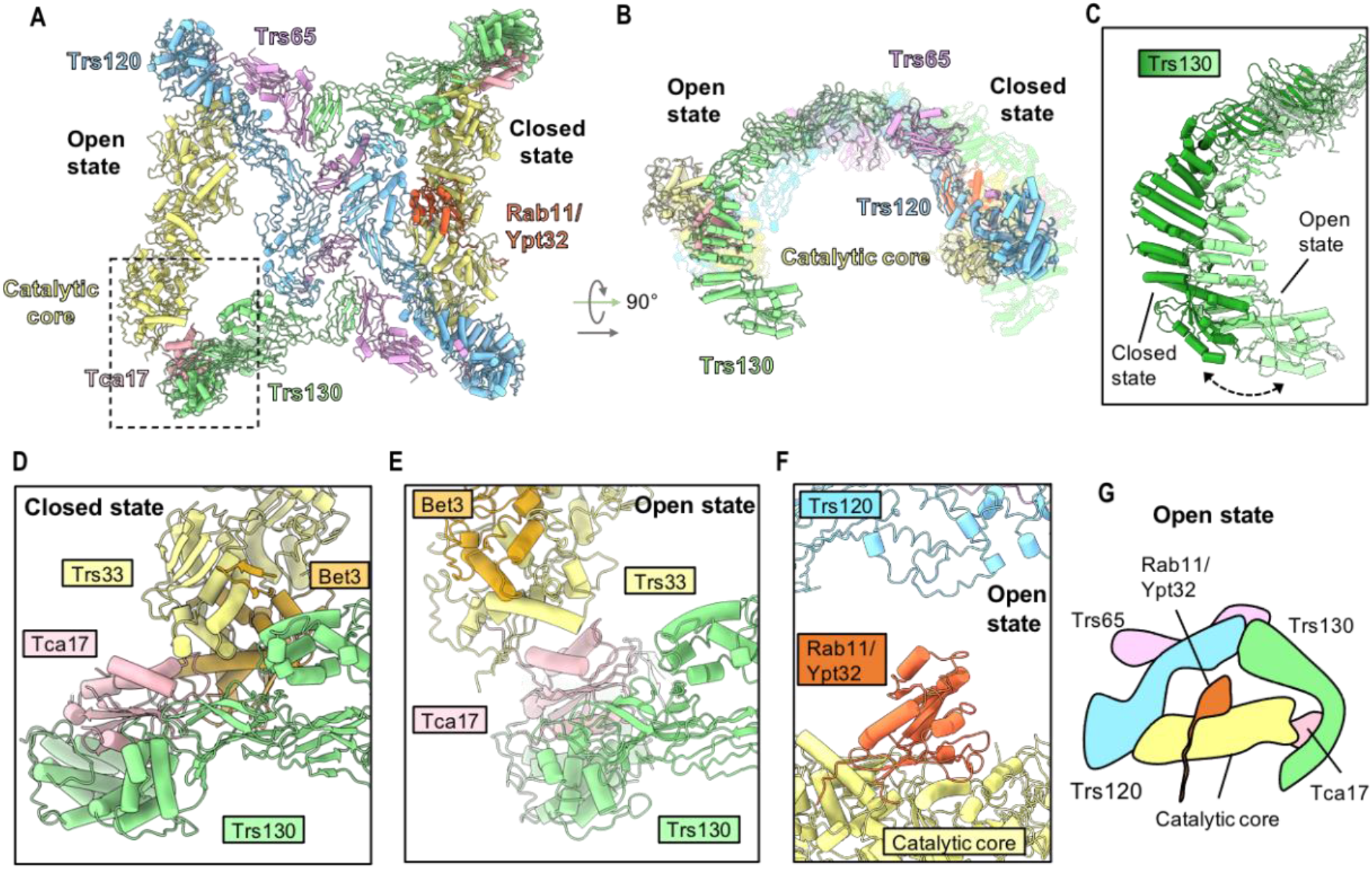
The open state of TRAPPII may facilitate access to the active site-chamber. (**A**) and (**B**) Structure of the closed/open state in which Rab11/Ypt32 is bound to the closed monomer. (**C**) Superposition of Trs130 from the closed and open states. (**D**) Close-up view of the interactions between Tca17 and the core in the closed state. (**E**) Same as (D) but for the open state. (**F**) Model of Rab11/Ypt32 superimposed onto the core of the open monomer to demonstrate the increased distance between the core and Trs120. (**G**) Schematic of the superimposed open state model in (F).

Compared to the closed state, in the open state the catalytic core has rotated 16 degrees, and the active site chamber has undergone significant expansion. This expansion is accomplished by two major structural changes. First, Trs130 has undergone a bending motion in which the leg region has moved more distal to its position in the closed state (Fig. 5C). Second, Tca17 adopts a completely different set of interactions with the core in the open and closed states. In the closed state Tca17 interacts with core subunits Bet3 and Trs33 (Fig. 5D and, fig. S9, A and B), while in the open state Tca17 has lost its interaction with Bet3 and interacts with a different surface of Trs33 (Fig. 5E and, fig. S9, C and D). Importantly, the Tca17-Bet3 interactions and each of the two different Tca17-Trs33 interactions involve conserved surfaces of both Bet3, Tca17, and Trs33 (although the surface of Tca17 involved in the open state interaction appears somewhat less conserved) (fig. S9, E-I). A mutation in TRAPPC6A, one of the two human paralogs of Trs33, is associated with a neurodevelopmental syndrome (*39*) and occurs at a position in the structure that appears important for stabilizing the open conformation (fig. S9D).

In the closed state, nucleotide-free Rab11 is tightly sandwiched in between the TRAPP core and the Trs120 lid, suggesting that conformational change is necessary to enable Rab11 to diffuse into or out of the TRAPPII active site. In the open state, the distance between the Trs120 lid and the TRAPP core has increased by ∼10 Å relative to the closed state (Fig. 5, F and G). This expansion of the active site chamber is expected to more easily accommodate initial binding of Rab11/Ypt32 to the catalytic core. We therefore propose that the open state facilitates Rab11 entrance to and exit from the active site chamber (Movie S1).

## Discussion

The structures presented here provide a structural basis for Rab11 activation by the TRAPPII complex, shed light on diseases associated with TRAPP genes, and explain the roles of each of the TRAPPII-specific subunits. Trs120 acts as a lid to enclose the active site, which is necessary for Rab11 activation because Rab11 is a relatively poor substrate for the TRAPP core. Trs130 provides a leg to lift the active site away from the membrane surface, enforcing counterselection against Rab1 by steric gating. Tca17 forms two different interfaces with Trs33, enabling the complex to adopt both closed and open conformations. The open state presumably allows Rab11 to enter and exit the active site chamber, while the closed state appears to be required for catalysis.

In the yeast TRAPPII complex, Trs65 creates an extensive interface to dimerize the complex. Metazoan TRAPPII complexes lack a homolog of the Trs65 subunit and are correspondingly monomeric (*26, 27, 30, 32, 40*). Dimerization may be important for enforcing the orientation of yeast TRAPPII on the membrane surface, so it is an open question whether the metazoan TRAPPII complex also employs steric gating to counterselect against Rab1. Nevertheless, comparison of our findings to published low-resolution structures, crosslinking data, and structural predictions of TRAPPII from other organisms (*26, 27, 29*) (figs. S5 and S6), indicates the overall organization of the TRAPPII monomer is conserved across eukaryotes. We therefore expect the mechanisms used by TRAPPII to identify and activate Rab11 to be conserved.

An additional factor may be involved in enforcing the orientation of TRAPPII on the membrane in both yeast and metazoans. Our group previously identified the GTPase Arf1 as a key recruiter of the TRAPPII (*21*), and we speculate that Arf1 may bind to a conserved surface on the bottom of the Trs120 subunit (fig. S5H). This would provide an additional “leg” for TRAPPII, opposite to the Trs130 leg. Support for this conjecture comes from biochemical reconstitution experiments in which Arf1 was required for enforcing TRAPPII substrate specificity towards Rab11 and against Rab1 (*23*). Future studies are needed to experimentally identify the Arf1 binding site on TRAPPII.

Our analysis of the structural data indicates that Tca17 plays a key role in conformational switching between the open and closed states of TRAPPII. However, although *TRS120* and *TRS130* are both essential genes, *TCA17* is not essential in yeast. Examination of the structures suggests that in the absence of Tca17, TRAPPII could still adopt both the open and closed states, but the complex would be more flexible and less stable. Although viable, cells lacking Tca17 appear to be significantly defective in activating Rab11 at the Golgi (*31*). Further analysis of the implications of switching between the open and closed states will require development of mutants that are trapped in either state.

More than one conformation was also observed for the fly TRAPPIII complex (*26*). This movement involved changes in the position of the TRAPPC8 subunit, which occupies a location in TRAPPIII similar to that of Trs120 in TRAPPII. The observed motion of TRAPPIII appears quite different from what we observe for TRAPPII. In one conformation TRAPPC8 was predicted to prevent Rab1 binding to the core, and in the other conformation TRAPPC8 was predicted to make direct contact with Rab1. Relative to the core binding site on the bottom face of the Rab, TRAPPC8 appeared to be positioned to interact with the side of Rab1, whereas we observe that the Trs120 lid interacts with the top of Rab11. Therefore, although TRAPPC8 was found to be important for Rab1 activation, based on the structural data it appears that TRAPPIII does not use TRAPPC8 as a lid (*26*). Additional studies are required to fully define the mechanisms used by metazoan TRAPPIII to distinguish and activate its substrate.

Rab GTPases interact with their effectors and regulators on the surface of organelle and vesicle membranes. The C-terminal HVD tails of Rabs create spacing between the effector-binding NBDs and the membrane surface. This spacing and flexibility is thought to be important for the function of Rab effectors such as motors and tethering factors. It is now clear that both the sequence and length of the HVD can also be critical determinants used by GEFs to identify their Rab substrates. Our work explains how a key Rab regulator meets its GTPase substrate where it is - at a distance from the membrane. Our findings support the idea that geometrical constraints imposed by organelle and vesicle membranes may be a general feature employed by proteins that interact on membrane surfaces to enforce specificity.

## Acknowledgments

We acknowledge the Cornell Center for Materials Research (CCMR), notably K. Spoth and M. Silvestry-Ramos, for access and support of electron microscopy sample preparation and data collection (NSF MRSEC program, DMR-1719875). We thank L. Thomas for valuable feedback on the manuscript and L. Thomas and R. Feathers for collecting preliminary cryo-EM data. This work was supported by NIH/NIGMS grant R35GM136258 to J.C.F.

## Author contributions

S.R.B. performed all experiments and data analysis. J.C.F. obtained funding and supervised the project. S.R.B. and J.C.F. wrote the manuscript.

## Competing interests

The authors declare no competing interests.

## Data and materials availability

All atomic coordinates and cryoEM maps will be deposited in the PDB and EMDB, respectively, and released upon publication.

## Materials and Methods

### Protein expression and purification

#### TRAPPII

Endogenous TRAPPII was purified from 48L of yeast with Trs130 tagged at the C-terminus with a tandem affinity purification (TAP) tag using the procedure described previously (*23*) and summarized here in brief. Yeast cells were homogenised using a freezer mill (SPEX SamplePrep) and the clarified cell lysate was applied to Sepharose 6B (Sigma-Aldrich) to remove protein that binds to Sepharose non-specifically. Sepharose 6B purified cell lysate was incubated with IgG Sepharose (GE Healthcare) followed by buffer wash steps to remove non-specifically binding proteins. TRAPPII bound to IgG resin was incubated with TEV protease overnight to remove the protein A-tag. IgG purified protein was further subjected to affinity purification using Calmodulin Sepharose (GE Healthcare). Purified TRAPPII was further subjected to size exclusion chromatography (SEC) using a Superdex 200 Increase 3.2/300 column (GE Healthcare). The final protein buffer was 10 mM Tris pH 8.0, 150 mM NaCl, 0.1% CHAPS, 1 mM Mg Acetate, 1 mM DTT. Peak elution fractions were pooled, concentrated and directly used for cryoEM analysis of the TRAPPII complex. An additional step was used to prepare the TRAPPII-Ypt32 complex as described below..

#### Rab11/Ypt32

A Rab11/Ypt32 construct harbouring a GST tag at the N-terminus was expressed recombinantly in Rosetta2 cells (EMD Millipore) and purified using the procedure described previously (*21*). The C-terminal cysteines (residues 221 and 222), which are prenylated in vivo, were substituted with a 7xHis tag in this construct. The final purified protein retained the C-terminal 7xHis tag while the GST tag was cleaved during the course of the purification.

### Preparation of TRAPPII-Rab11/Ypt32 complex

Ten-fold molar excess of purified Rab11/Ypt32-7xHis was added to the TEV cleavage reaction mixture in the TRAPPII purification procedure described above. Calf intestine alkaline phosphatase (Invitrogen) was added to the reaction mixture and incubated at 4°C overnight to hydrolyze the nucleotide and facilitate stable association of nucleotide-free Rab11/Ypt32 with TRAPPII. The TRAPPII-Rab11/Ypt32 complex was further purified using the affinity chromatography with Calmodulin Sepharose and SEC as described above.

### CryoEM sample preparation, data collection, data processing and model building

Cryo-EM grids for TRAPPII and TRAPPII-Rab11/Ypt32 complexes were prepared using the same procedure. 3μL sample (TRAPPII at 6.8mg/ml and TRAPPII-Rab11/Ypt32 at 5.5mg/ml) was applied to a plasma cleaned UltrAuFoil R1.2/1.3 (Quantifoil) grid inside a Vitrobot IV (Thermo Fisher Scientific) at 4 °C and 100% humidity. The sample was incubated on the grid for 10 seconds, followed by blotting for 5 seconds using blot force 3, and then immediately plunged into liquid nitrogen-cooled liquid ethane.

Cryo-EM data were collected using a Talos Arctica operating at 200kV and equipped with a K3 detector and BioQuantum energy filter. For the TRAPPII-Rab11/Ypt32 data set, 4,998 50-frame movies were collected in super resolution mode (0.62Å/super resolution pixel). Frame alignment was carried out using MotionCor2 (*41*) and initial defocus estimation was carried out using Patch CTF in cryoSPARC (*42*). 4,906 micrographs showing CTF resolution estimates higher than 6Å were selected for further processing. The first 200 micrographs were used to pick particles using a reference-free particle picker (“Blob picker”). These particles were subjected to 2D classification and classes showing sharp protein-like features were selected as templates for reference based particle picking (“Template picker”) to obtain an initial set of 979,187 particles which were fourier cropped to 1.43Å/pixel. Additional steps of 2D classification and two class ab-initio refinement followed by a two class heterogenous refinement were carried out to remove junk particles which gave a clean set of 524,578 particles. This clean particle set was then imported into RELION 3.1 (*43, 44*) and subjected to iterative rounds of 3D refinement with C2 symmetry and CTF refinement followed by a Bayesian polishing step to obtain a consensus 3D reconstruction at 3.9Å. In order to assess the heterogeneity of the data set, unmasked 3D classification was carried out using C1 symmetry which revealed multiple conformational states of the TRAPPII-Rab11/Ypt32 complex as summarized in fig. S2. We observed three distinct classes for the TRAPPII-Rab11/Ypt32 complex: one in which both the monomers were in the “closed” state (“closed/closed” dimer); one in which one monomer was in the “closed” while the other monomer was in the “open” state (“closed/open” dimer); and a third state in which one monomer was in the “closed” state and the other monomer was in a “partially open” state (“closed/partially open” dimer). The particles in the “closed/closed” dimer class were then subjected to 3D refinement using C2 symmetry to obtain a 4.1Å final reconstruction of the symmetric dimer. In this map, the local resolution of the central region was higher than that of the regions furthest from the central region. We suspected this could be due to relative movement between the two monomers and therefore we carried out symmetry expansion of the particle set in which each particle image was superposed on itself after applying a rotation of 180°. We then subtracted the signal for one of the monomers from this symmetry expanded data set and carried out 3D refinement on the subtracted particle images to obtain a reconstruction of the closed monomer. Further 3D classification was used to select for Rab11/Ypt32 bound closed monomers to obtain a final reconstruction of the closed Rab11/Ypt32-bound monomer at 3.7Å.

The particles in the “closed/open” and “closed/partially open” states were combined and subjected to 3D refinement to obtain a consensus map of the asymmetric dimer. These particles were then subjected to 3D classification with fixed alignment which gave two “closed/partially open” classes and two “closed/open” classes. The classes with a “partially open” monomer showed loss of map density at the interface between Trs130, Tca17 and the catalytic core, possibly due to high conformational heterogeneity of this region. Among the two “closed/open” classes, one class showed map density for Ypt32 bound to the “closed” monomer and this class was further refined to obtain a 4.5Å resolution reconstruction (asymmetric dimer “state A”). The other “closed/open” class refined to 4.8Å (asymmetric dimer “state B”). We then subtracted the signal for the “closed” monomer from both the “closed/open” dimer classes to obtain focused maps for the “open” monomers from these classes. To further improve the resolution, the signal subtracted particles of “open” monomers from both the classes were combined to obtain a final reconstruction of the “open” monomer at an overall resolution of 4.2Å. A 4.2Å reconstruction for the “closed”-Rab11/Ypt32 bound monomer from asymmetric dimer state A was obtained upon subtraction of signal for the “open” monomer. Significant improvements in resolution for portions of the “open” and “closed” monomers and the symmetric and asymmetric dimer maps were obtained by doing more focused refinements as shown in fig. S2.

For the TRAPPII-only data set, 3,333 50-frame movies were collected in super resolution mode (0.62Å/super resolution pixel). A similar procedure to that described for the TRAPPII-Rab11/Ypt32 data set was used to obtain a clean set of 303,062 particles (1.43Å/pixel). Unlike the TRAPPII-Rab11/Ypt32 dataset, all further processing steps were carried out in cryoSPARC (fig. S3). These particles were further subjected to 3D classification (heterogeneous refinement) which yielded four distinct populations of TRAPPII - a “closed/open” state, a “closed/closed” state, a “partially open/open” state and a “closed/partially open” state. Particles in each of these states were subjected to homogenous 3D refinement followed by non-uniform 3D refinement to obtain reconstructions of the closed/open, closed/closed, partially open/open, closed/partially open states at 4.7Å, 4.2Å, 4.9Å and 4.9Å, respectively.

### Model building and refinement

The symmetric dimer map, the monomer map and the corresponding focused maps were used for building the symmetric TRAPPII-Rab11/Ypt32 model. For the asymmetric TRAPPII-Rab11/Ypt32 model, the asymmetric dimer state A map, the map for the closed monomer from asymmetric dimer state A, the open monomer map, and the corresponding focused maps were used. Density modification (*45*) was performed on these maps in Phenix (*46*) to facilitate interpretation during model building. Individual models for the catalytic core subunits Trs23, Trs31, Bet5, Bet3 from PDB 3CUE, the predicted models for Trs20 and Trs33 obtained using trRosetta, the model for Rab11/Ypt32 from PDB 3RWO and the model for Tca17 from PDB 3PR6 were fitted into the focused maps for the core/Ypt32 region in Chimera (*47*) and rebuilt in Coot (*48*). Predicted models for Trs120, Trs130 and Trs65 were initially obtained using trRosetta(*49*). The predicted models for Trs120, Trs130 and Trs65 did not fit well in the cryo-EM maps due to differences in the overall organization of domains in the predicted models versus that observed in our cryoEM maps. Hence, the predicted models for Trs120, Trs130 and Trs65 were split into individual domains that fit well into the focused maps and were further rebuilt in Coot. Notably, the low local resolution of map density for parts of the N-terminal domains of Trs65 (residues 1-120 and 198-210) and Trs130 (1-239) precluded accurate side chain modelling and therefore we built these regions as poly-alanine models. A model for the closed and open monomers was made by fitting the rebuilt models for the individual subunits and domains into the closed and open monomer maps respectively. These composite monomer models were further rebuilt into the respective monomer maps to ensure residues at the focused map interfaces were built correctly. Composite models for the symmetric (closed/closed) and the asymmetric (closed/open) dimers were made by fitting the respective monomer models into the consensus maps for the symmetric and asymmetric dimers respectively. The overall models were then subjected to real-space refinement against composite maps for the symmetric (closed/closed) and the asymmetric (closed/open) dimers. The composite maps were generated with Combine Focused Maps in Phenix (*46*) using the density-modified focused maps together with the monomer and dimer consensus maps. Real-space refinement was carried out in Phenix using secondary structure and Ramachandran restraints (*50*). Model-map validation statistics of the refined models were calculated using the Comprehensive validation tool in Phenix (*50*). The 0.143 model-map FSCs are reported in fig. S4 and cryo-EM data collection, refinement, and model validation statistics are reported in Table 1.

### Software

The structural biology software we used is maintained by SBGrid (*51*).

### Fluorescence Microscopy

Cells were grown in appropriate synthetic dropout media at 30°C to mid-log phase (OD600 of ∼0.5) and imaged using DeltaVision Elite system (GE Healthcare Life Sciences) equipped with an Olympus IX-71 inverted microscope, DV Elite complementary metal-oxide semiconductor camera, a 100×/1.4 NA oil objective, and a DV Light SSI 7 Color illumination system with Live Cell Speed Option with DV Elite filter sets. Images were acquired and deconvoled (conservative setting, six cycles) using DeltaVision software softWoRx 6.5.2 (Applied Precision). All fluorescence microscopy images shown in figure panels are single focal planes.

### GEF-Rab Interaction Test (GRab-IT)

GRab-IT assays were performed as described previously (*23*). N-terminal mRFPmars-tagged Rab baits were expressed as nucleotide-free mutant constructs (Rab1/Ypt1(D124N) and Rab11/Ypt31/32(D129N)) and were ectopically localized to the mitochondrial membrane by substituting the C-terminal cysteine residues with the transmembrane domain of Fis1 (residues 129-155). The endogenous copy of the GEF (Trs130 or Trs85) was tagged with a C-terminal mNeonGreen tag. Recruitment of the GEF on Rab constructs localized to the mitochondria was quantified using Manders overlap analysis in ImageJ (JACoP plugin). Images were cropped to contain three-to-five cells and Manders overlap coefficient was calculated for ≥ 30 cells. Statistical significance was determined using a one-way ANOVA (ANalysis Of VAriance) with post-hoc Tukey’s HSD (Honestly Significant Difference) for multiple comparisons (*p < 0.05, **p < 0.01, ***p < 0.001).

### Complementation Tests

Plasmid shuffling assays were performed to test if Rab11/Ypt32 mutants support growth in the absence of wild type gene using the procedure described previously (*23*). A similar procedure was used to test Trs120 mutants. *ypt31*Δ*ypt32*Δ and *trs120*Δ null mutant yeast were maintained by a copy of *YPT32* and *TRS120* respectively, expressed using the pRS416 vector (URA selection). These strains were transformed with pRS415 vectors (LEU selection) containing the mutant and wild type versions of *YPT32* and *TRS120*. Transformed cells were serially diluted and grown on synthetic dropout media supplemented with 3.9 mM 5-fluoroorotic acid (5-FOA) at 30°C.

### Sequence conservation analysis

Sequence alignments for Rab1 and Rab11 homologs were performed using Clustal Omega. Consurf analysis (*52*) was performed using the default settings for Trs120, Trs130 and Trs65 for closely related homologs. For Consurf analysis of Rab11/Ypt32, custom sequence alignment of Rab11 homologs across the model organisms described above was provided to prevent contamination with non-Rab11 protein sequences.

**Table S1.** CryoEM data collection and model validation statistics.

|  | Symmetric dimer<br>(composite structure)<br>(EMDB-xxxx)<br>(PDB xxxx) | Symmetric dimer<br>(consensus map)<br>(EMDB-xxxx) | Closed monomer<br>(EMDB-xxxx) | Asymmetric dimer state A<br>(composite structure)<br>(EMDB-xxxx)<br>(PDB xxxx) | Asymmetric dimer state A (consensus map) (EMDB-xxxx) | Open monomer<br>(EMDB-xxxx) | Closed monomer from asymmetric dimer state A<br>(EMDB-xxxx) |
| --- | --- | --- | --- | --- | --- | --- | --- |
| **Data collection and processing** |  |  |  |  |  |  |  |
| Nominal Magnification | 63000 |  |  |  |  |  |  |
| Voltage (kV) | 200 |  |  |  |  |  |  |
| Electron exposure<br>(e-/Å²) | 53.2 |  |  |  |  |  |  |
| Defocus range (µm) | -0.8 to -2.5 |  |  |  |  |  |  |
| Pixel size (Å) | 1.24 |  |  |  |  |  |  |
| Symmetry imposed | C2 | C2 | C1 | C1 | C1 | C1 | C1 |
| Initial particle images (no.) | 524,578 | 524,578 | 524,578 | 524,578 | 524,578 | 524,578 | 524,578 |
| Final particle images (no.) | NA | 163,758 | 369,488 | NA | 74,313 | 149,906 | 74,313 |
| Map resolution (Å)<br>FSC threshold (0.143) | NA | 4.1 | 3.7 | NA | 4.5 | 4.2 | 4.2 |
| Map resolution range (Å) | NA | 3.7 - 12.9 | 3.5 - 9.7 | NA | 4 - 15.1 | 4 - 10.7 | 4 - 11.5 |
| **Model Refinement** |  |  |  |  |  |  |  |
| Initial model used<br>(PDB code) | 3CUE, 3RWO,<br>3PR6 |  |  | 3CUE, 3RWO,<br>3PR6 |  |  |  |
| Model resolution (Å)<br>FSC threshold (0.5) | 3.5 |  |  | 3.9 |  |  |  |
| Map sharpening *B*<br>factor (Å²) | NA | -94 | -81 | NA | -92 | -77 | -87 |
| Model composition<br>Non-hydrogen atoms<br>Protein residues<br>Ligands | 62838<br>7798<br>PLM: 4 |  |  | 60,829<br>7507<br>PLM: 4 |  |  |  |
| **B factors (Å<sup>2</sup>)**<br>Protein<br>Ligand | 100.04<br>85.22 |  |  | 104.69<br>81.62 |  |  |  |
| **R.m.s. deviations**<br>Bond lengths (Å)<br>Bond angles (°) | 0.004<br>0.838 |  |  | 0.005<br>0.886 |  |  |  |
| **Validation**<br>MolProbity score<br>Clashscore<br>Poor rotamers (%) | 1.73<br>4.10<br>0.04 |  |  | 1.86<br>5.17<br>0.16 |  |  |  |
| **Ramachandran plot**<br>Favored (%)<br>Allowed (%)<br>Disallowed (%) | 90.55<br>9.34<br>0.12 |  |  | 88.78<br>10.83<br>0.39 |  |  |  |

**Fig. S1.**
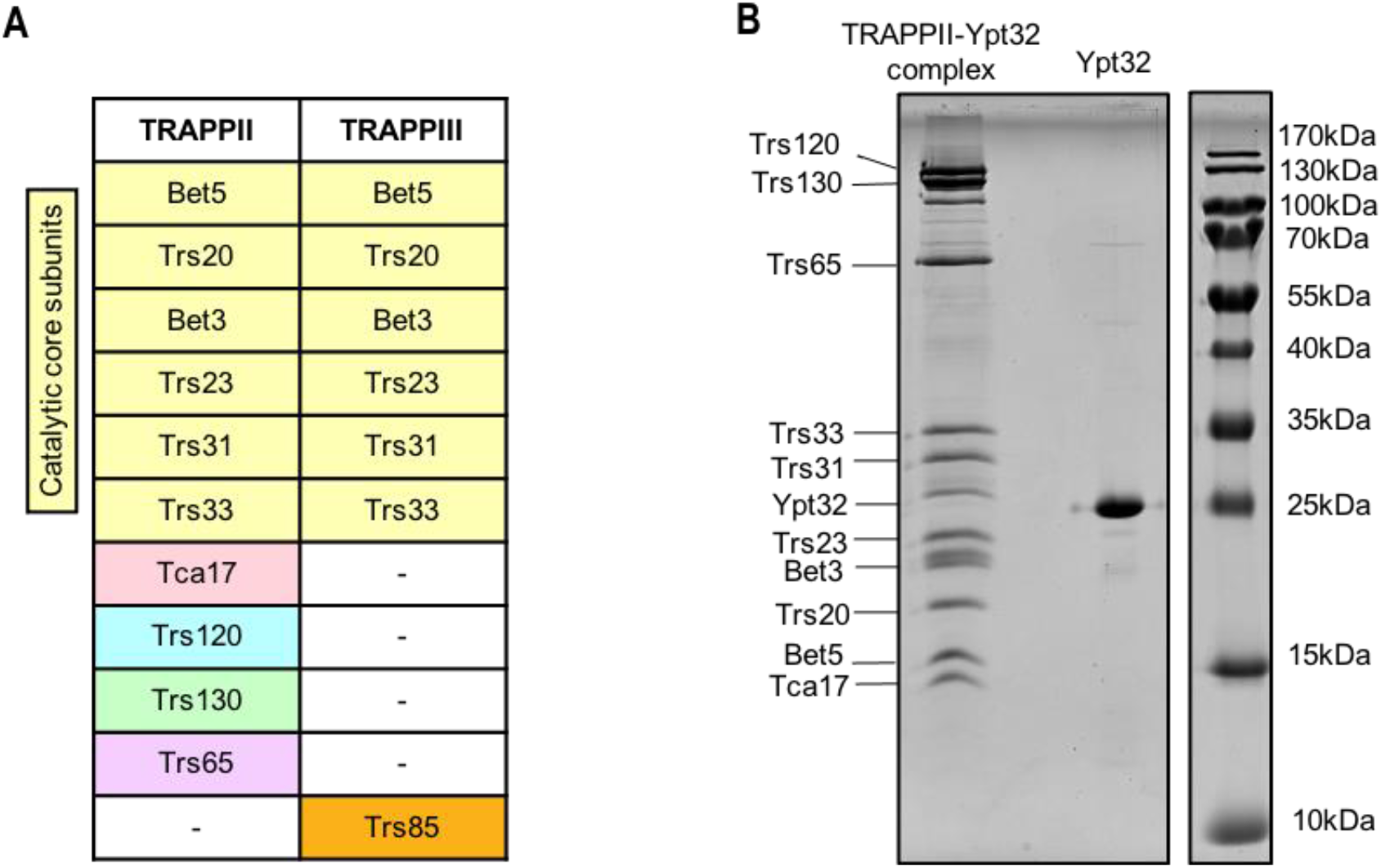
CryoEM sample preparation. **(A)** Table detailing the subunit composition of the yeast TRAPPII and TRAPPIII complexes. **(B)** SDS-PAGE gel showing purified TRAPPII-Rab11/Ypt32 complex and purified Rab11/Ypt32.

**Fig. S2.**
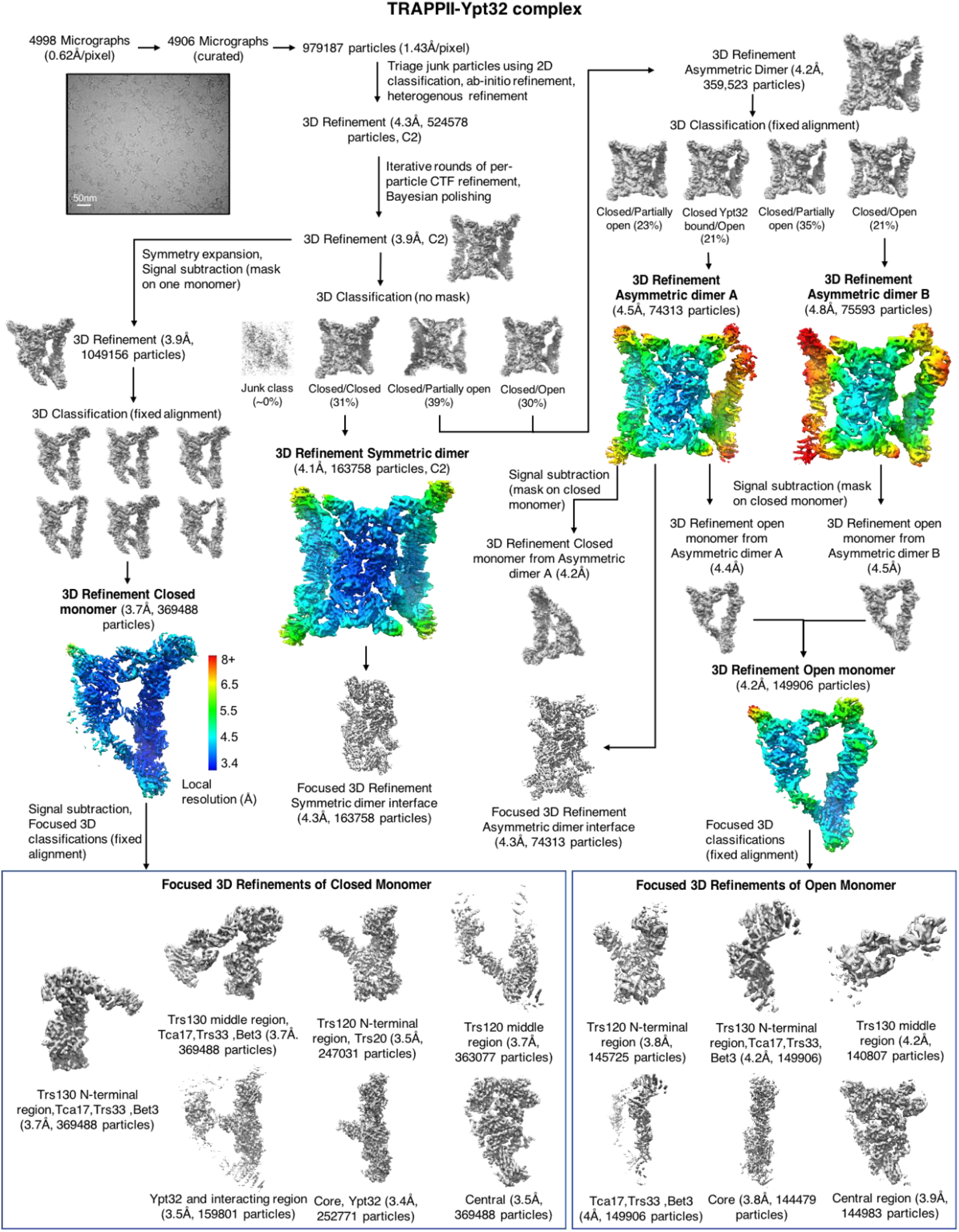
CryoEM data processing pipeline for TRAPPII-Rab11/Ypt32 complex. Flowchart illustrating the data processing strategy for the TRAPPII-Rab11/Ypt32 complex cryoEM data (see Methods).

**Fig. S3.**
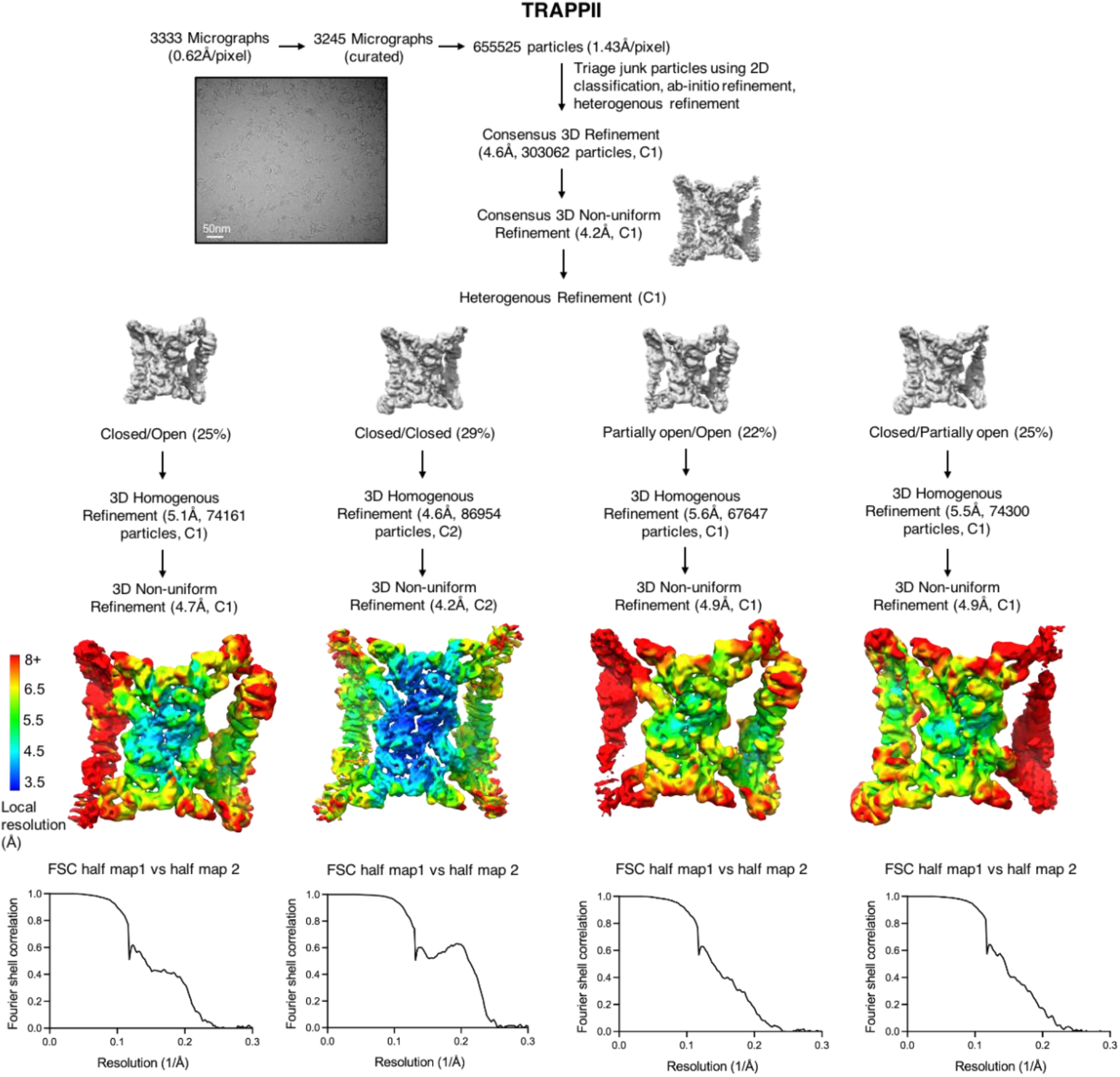
CryoEM data processing pipeline for TRAPPII. Flowchart illustrating the data processing strategy for the TRAPPII-only cryoEM data (see Methods). Fourier shell correlation plots for the indicated refinement reconstructions are shown below the maps.

**Fig. S4.**
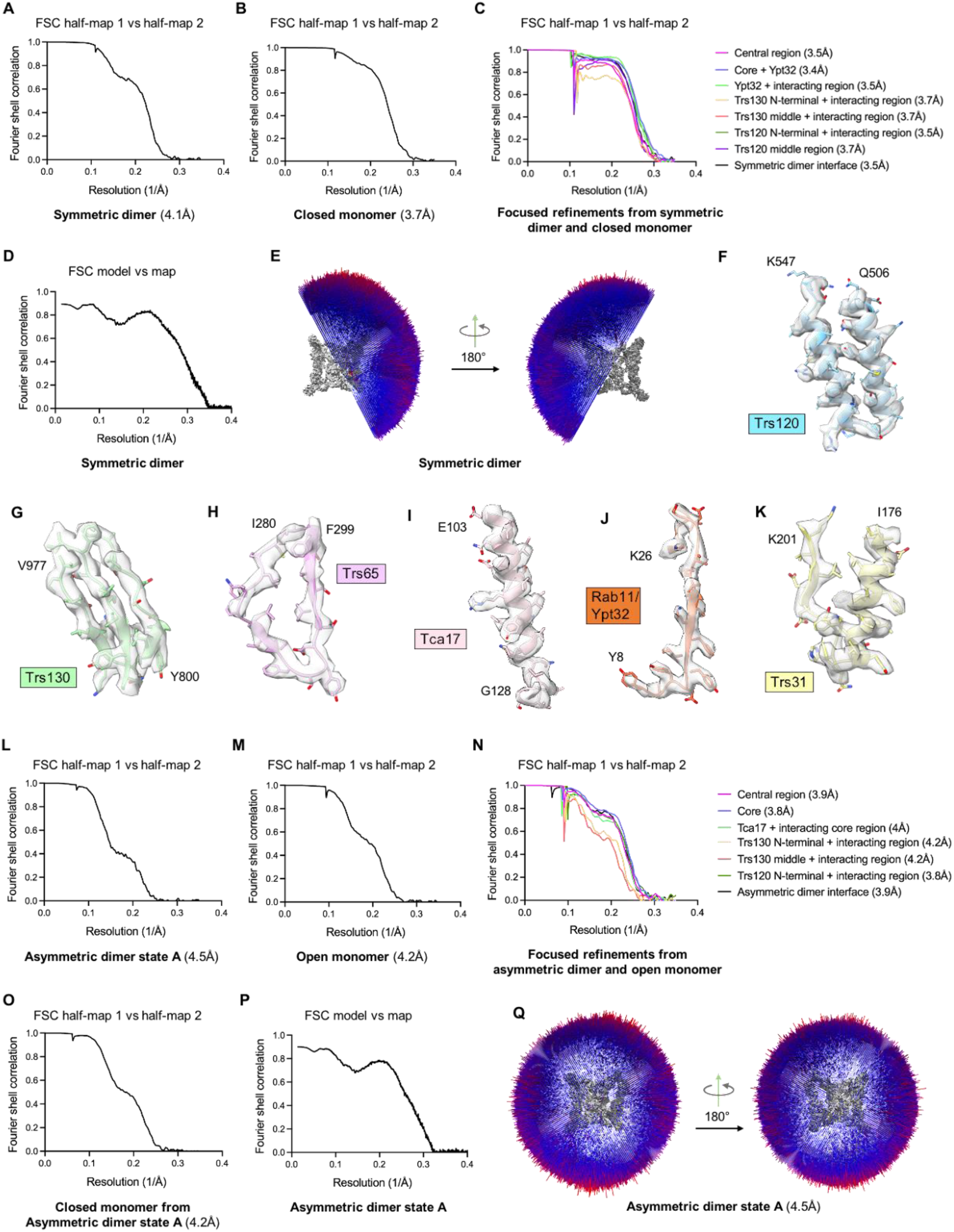
CryoEM map quality and model-map fit. (**A**) Fourier shell correlation plot for the symmetric (closed/closed) TRAPPII-Rab11/Ypt32 dimer reconstruction. (**B**) Fourier shell correlation plot for the symmetry-expanded TRAPPII-Rab11/Ypt32 closed monomer reconstruction. (**C**) Fourier shell correlation plots for the indicated focused refinement reconstructions produced from the symmetric TRAPPII-Rab11/Ypt32 complex. (**D**) Fourier shell correlation plot comparing the refined symmetric TRAPPII-Rab11/Ypt32 dimer model to the symmetric (closed/closed) TRAPPII-Rab11/Ypt32 dimer reconstruction. (**E**) Orientation histogram for the symmetric (closed/closed) TRAPPII-Rab11/Ypt32 dimer particle refinement. (**F**) Example cryoEM density for the Trs120 subunit. (**G**) Example cryoEM density for the Trs130 subunit. (**H**) Example cryoEM density for the Trs165 subunit. (**I**) Example cryoEM density for the Tca17 subunit. (**J**) Example cryoEM density for Rab11/Ypt32. (**K**) Example cryoEM density for the Trs31 subunit (as an example of the core complex). **(L)** Fourier shell correlation plot for the asymmetric (closed/open) dimer “state A” reconstruction. **(M)** Fourier shell correlation plot for the open TRAPPII monomer reconstruction. (**N**) Fourier shell correlation plots for the indicated focused refinement reconstructions produced from the open monomer. (**O**) Fourier shell correlation plot for the closed TRAPPII monomer reconstruction from the asymmetric (closed/open) dimer “state A”. (**P**) Fourier shell correlation plot comparing the refined symmetric asymmetric dimer model to the asymmetric dimer reconstruction. (**Q**) Orientation histogram for the asymmetric dimer “state A” particle refinement.

**Fig. S5.**
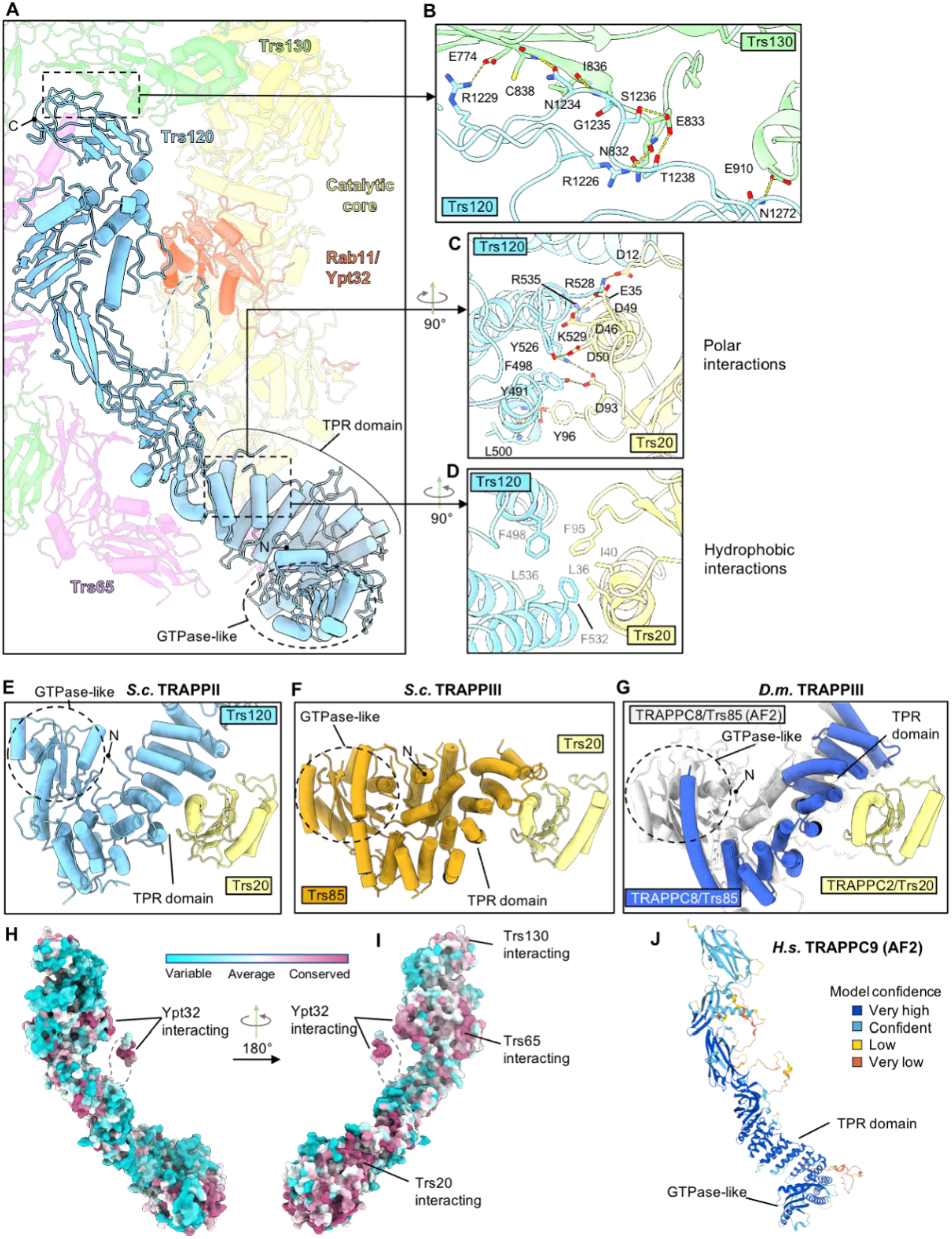
Analysis of Trs120 structure and interactions within the TRAPPII complex. (**A**) Overall fold of Trs120. TPR : tetratricopeptide repeat. (**B**) Interactions at the Trs120-Trs130 interface. (**C**) Polar interactions at the Trs120-Trs20 interface. (**D**) Hydrophobic interactions at the Trs120-Trs20 interface. (**E-G**) Comparison of Trs120-Trs20 in TRAPPII (E, this work), Trs85-Trs20 in yeast TRAPPIII (*25*) (F), and TRAPPC8-TRAPPC2 in fly TRAPPIII(*26*) (G) interfaces. Note that the GTPase-like domain of TRAPPC8 was not modeled in the published fly TRAPPIII structure, so we superimposed the AlphaFold prediction (*53*) (shown in gray) onto the fly TRAPPC8 experimental model (*26*). (**H**,**I**) Conservation analysis of Trs120. (**J**) AlphaFold prediction of the human TRAPPC9 structure (*53*) shows an overall fold similar to that of Trs120.

**Fig. S6.**
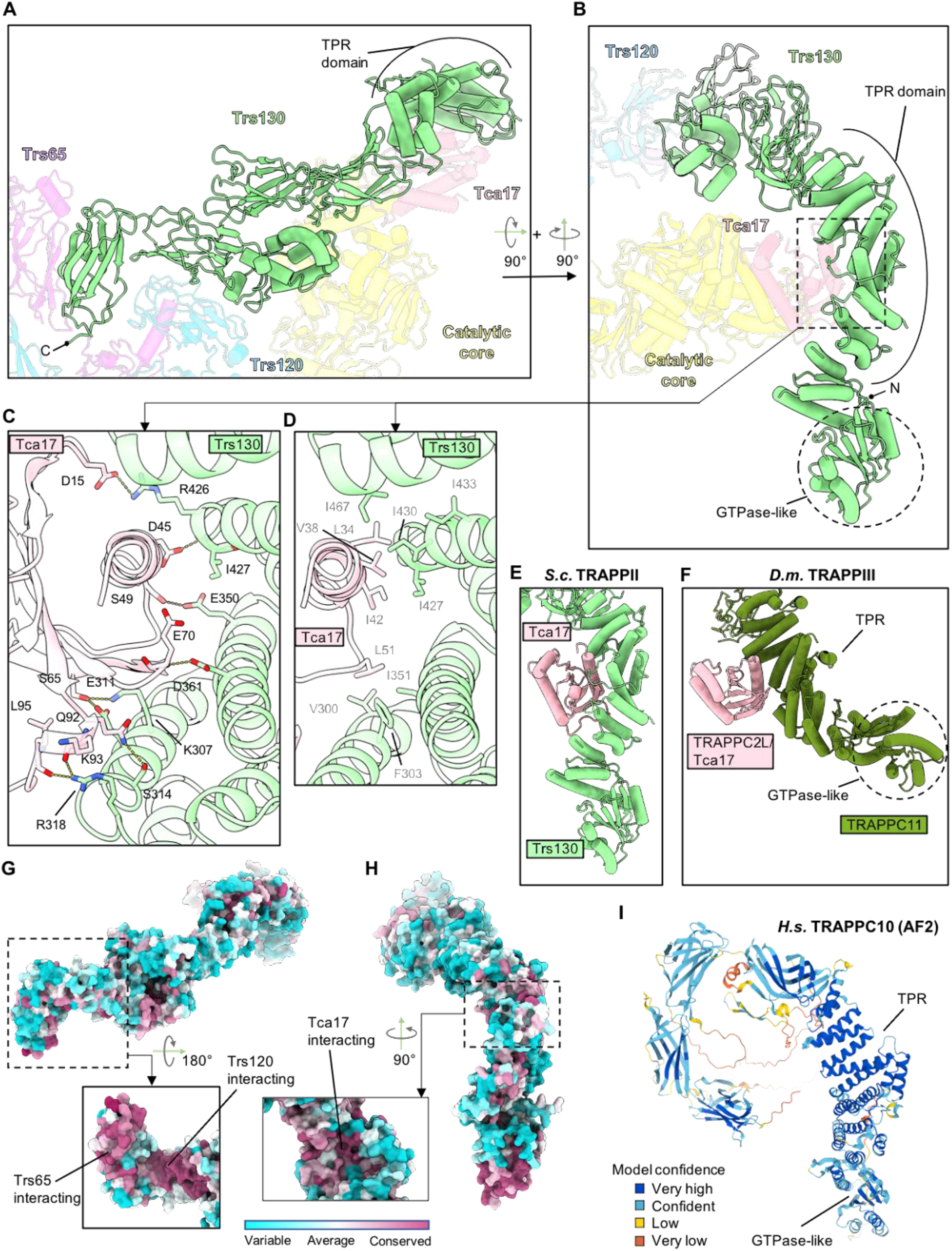
Analysis of Trs130 structure and interactions within the TRAPPII complex. (**A**,**B**) Overall fold of Trs130. TPR : tetratricopeptide repeat. (**C**) Polar interactions at the Trs130-Tca17 interface. Tca17 residue D45 is equivalent to a mutation in human TRAPPC2L associated with a neurodevelopmental disorder (*36*) (**D**) Hydrophobic interactions at the Trs130-Tca17 interface. (**E**,**F**) Comparison of the leg elements from Trs130 (E, this work) and TRAPPC11 from fly TRAPPIII (*26*) (F). Note that the leg of TRAPPC11 from fly TRAPPIII appears “bent” relative to the “straight” leg of Trs130. (**G**,**H**) Conservation analysis of Trs130. (**I**) AlphaFold structure prediction (*53*) of the human TRAPPII subunit TRAPPC10 (Trs130 paralog) shows an overall fold similar to that of Trs130. Note that the TRAPPC10 leg appears “straight”, similar to the structure of yeast Trs130 but unlike the corresponding region of TRAPPC11 which appears “bent” in fly TRAPPIII.

**Fig. S7.**
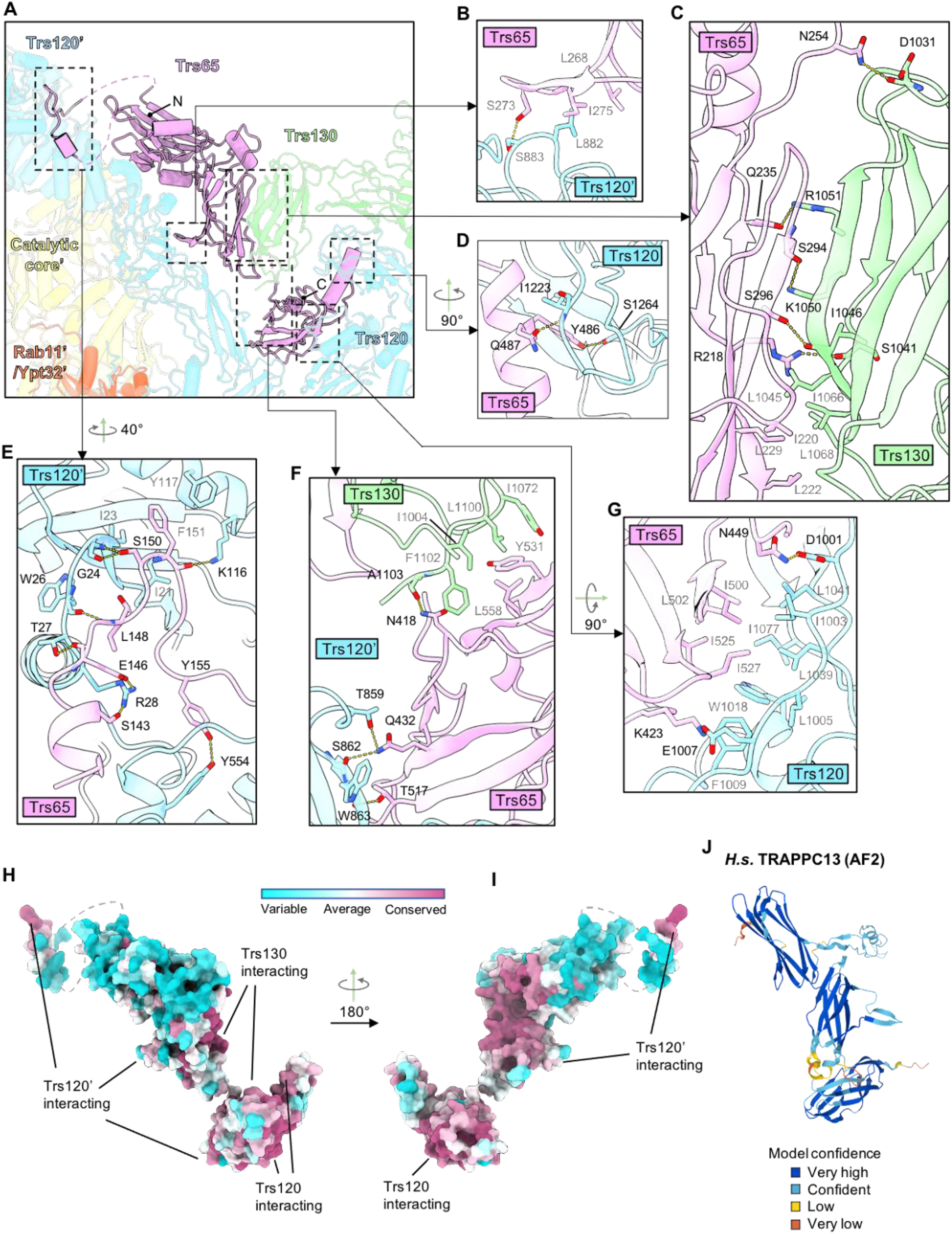
Analysis of Trs65 structure and interactions within the TRAPPII complex. (**A**) Overall fold of Trs65. Labels with prime (‘) symbols indicate subunits belonging to the other monomer within the TRAPPII dimer. (**B-G**) Interactions between Trs65 and other subunits. (**H**,**I**) Conservation analysis of Trs65. (**J**) AlphaFold structure prediction (*53*) of human TRAPPC13 shows an overall fold similar to that of Trs65. Note that although it exhibits structural similarity to Trs65, TRAPPC13 is part of the metazoan TRAPPIII complex (*30, 40*).

**Fig. S8.**
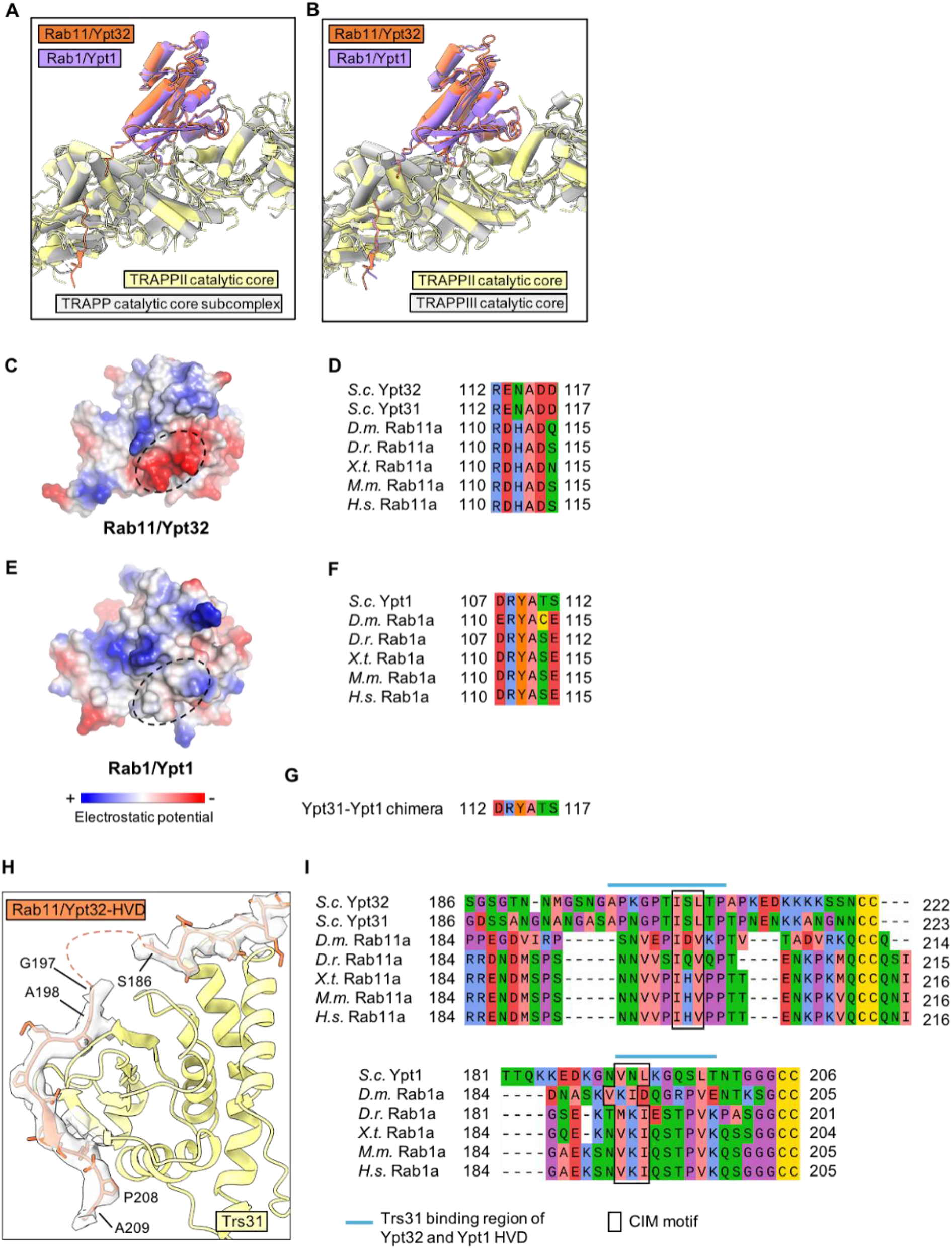
Differences between Rab1 and Rab11 TRAPP core binding interactions. (**A**) Superposition comparing the structure of TRAPPII-bound Rab11/Ypt32 (this study) to TRAPP core-bound Rab1/Ypt1 (*22*). (**B**) Superposition comparing the structure of TRAPPII-bound Rab11/Ypt32 (this study) to TRAPPIII bound Rab1/Ypt1 (*25*). (**C-F**) Comparison of the surfaces of Rab11/Ypt32 and Rab1/Ypt1 to highlight the electrostatic differences (**C**,**E**) of sequences (**D**,**F**) located near the TRAPP-core binding site of each Rab. (**G**) Sequence of the region used in the Ypt31-Ypt1 graft chimera construct used for experiments shown in Fig. 2, F and G. (**H**) Close-up of the interaction between the Rab11/Ypt32 HVD (orange) and the Trs31 core subunit in TRAPPII (yellow). CryoEM density of the HVD is shown in white with black outline. (**I**) Alignments of the Rab1 and Rab11 HVD regions are aligned to each other using the Rab1/Ypt1 and Rab11/Ypt32 binding-sites on Trs31 (blue line) as a structural alignment reference. The CIM motifs required for prenylation are indicated.

**Fig. S9.**
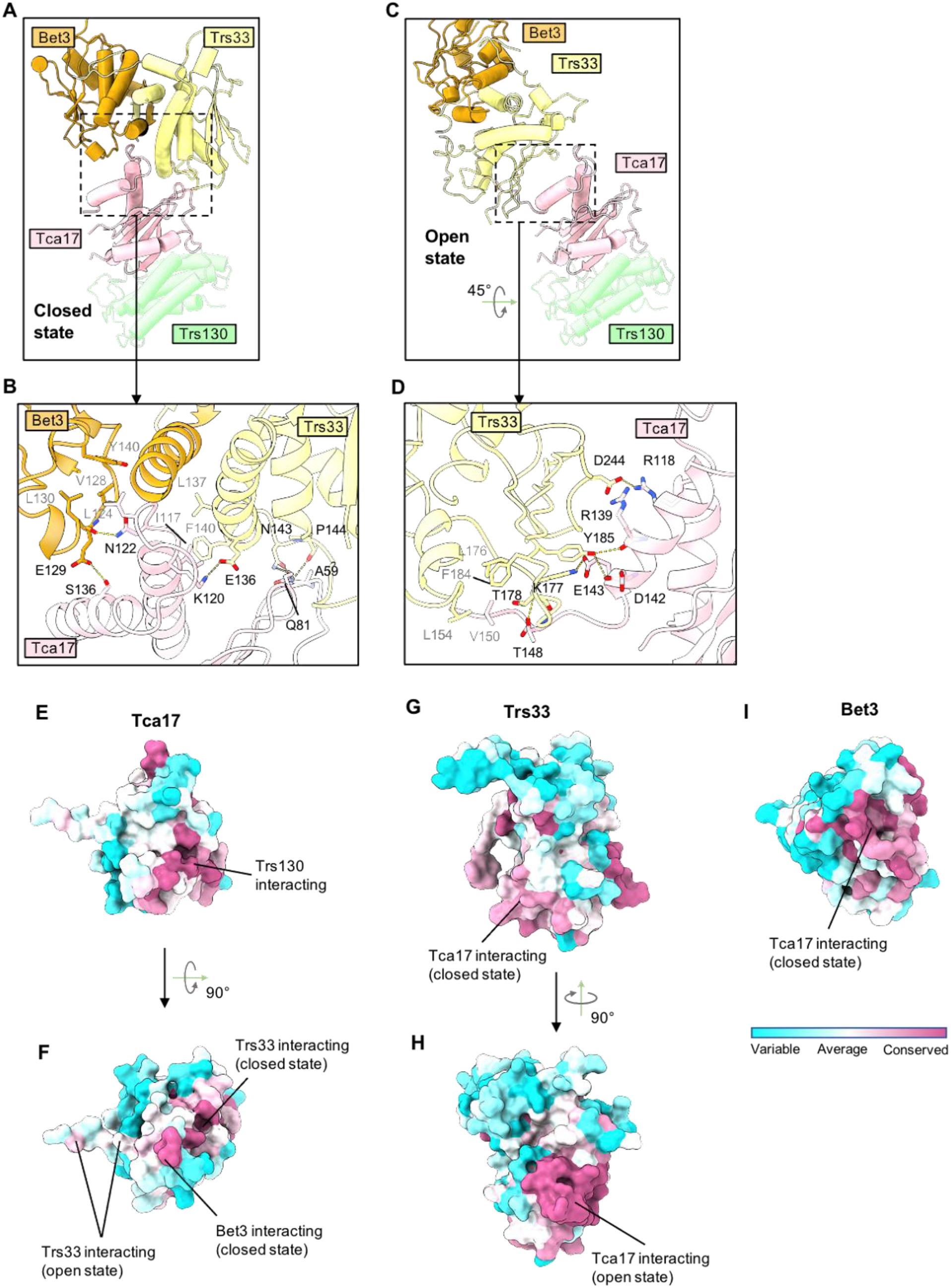
Distinct interactions between Tca17 and the core in the open and closed states. (**A**,**B**) Close-up view of the interactions between Tca17 and the core subunits Trs33 and Bet3 in the closed state. (**C**,**D**) Close-up view of the interactions between Tca17 and Trs33 in the open state. Note that Trs33 F184 corresponds to the residue position of a substitution mutation in human TRAPPC6A that is associated with a neurodevelopmental syndrome (*39*) (the disease allele results in a TRAPPC6A Y93N substitution according to the UNIPROT database sequence of the human protein). (**E**,**F**) Conservation analysis of Tca17, indicating the interaction sites. (**G**,**H**) Conservation analysis of Trs33. (**I**) Conservation analysis of Bet3.

**Fig. S10.**
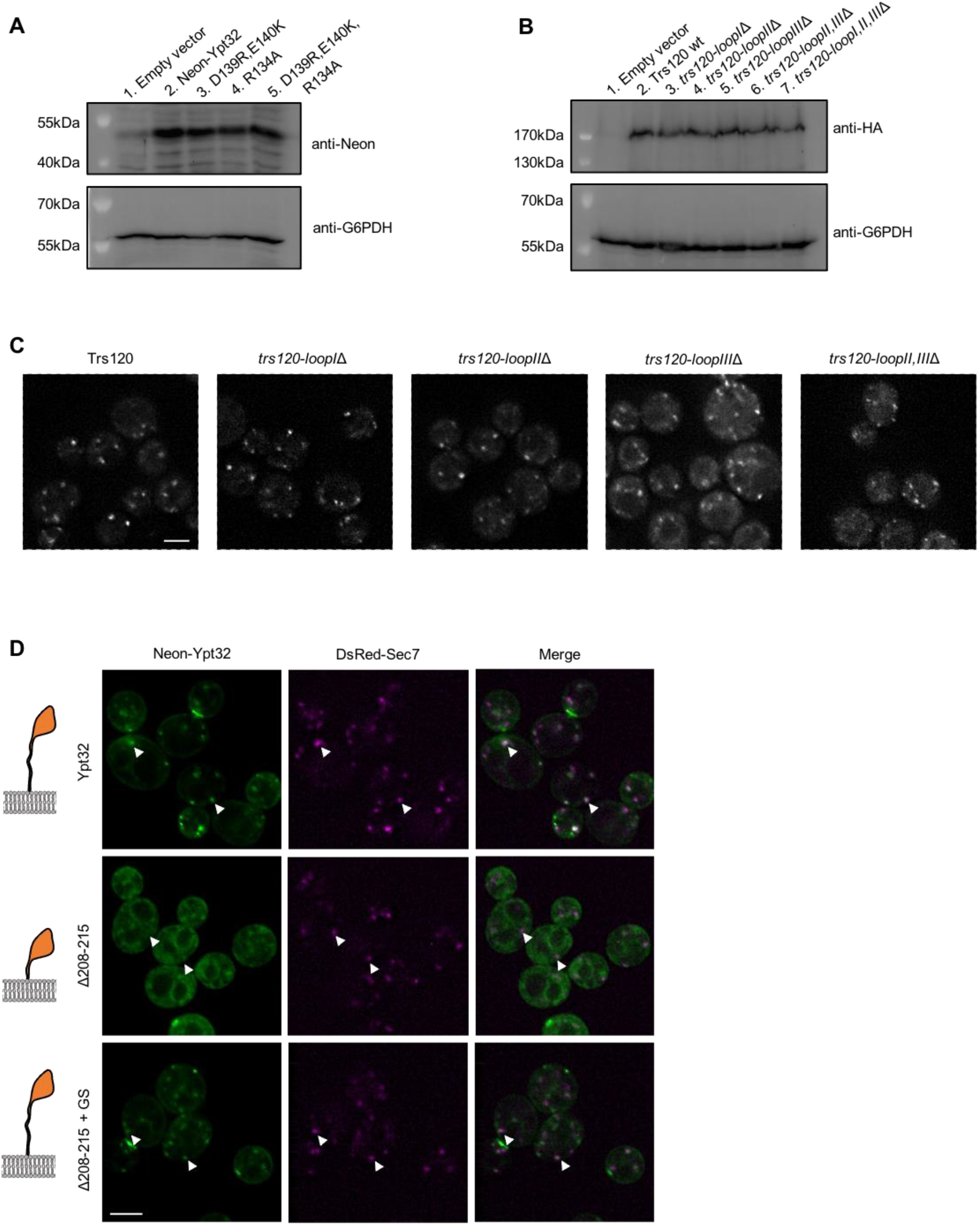
Western blot analysis of Rab11/Ypt32 mutants and localization of Trs120 and Rab11/Ypt32 mutants. (**A**) Immunoblot to confirm expression of the mutants shown in Fig. 3E. (**B**) Immunoblot to confirm expression of the HA-tagged Trs120 and loop mutants shown in Fig. 3F. (**C**) Imaging data using Trs120-mNeonGreen constructs indicating that the Trs120 loop deletion constructs are expressed and localized similar to the wild-type. (**D**) Localization of an extra copy of wild-type and mutant mNeonGreen-Rab11/Ypt32 constructs in yeast. Colocalization with Sec7-marked late-Golgi compartments is indicative of active Rab11/Ypt32. Scale bars shown in (C) and (D) are 2µm.

**Movie S1. Mechanism of activation of Rab11/Ypt32 by the TRAPPII complex**.

Coloring is the same as in Fig. 1. First, the architecture of the complex is presented, then the proposed orientation of the complex on the membrane is shown. The “lid” and “leg” features are then highlighted. Next, two conformations adopted by the complex are shown. Based on the structural and functional results, we then present the structural changes the complex likely undergoes during the exchange mechanism: the TRAPPII monomer in the closed conformation is viewed from the side, the movie then shows a “morph” transition to the open conformation. In the next stage, GDP-bound Rab11/Ypt32 diffuses into the active site chamber and binds to the catalytic core. The structure then transitions back to the closed confirmation, triggering GDP release and conformational change of Rab11/Ypt32. GTP then binds to Rab11/Ypt32, as the concentration of GTP is significantly higher than the concentration of GDP within cells. GTP-binding triggers another conformational change of Rab11/Ypt32 to its active state. This conformation of Rab11/Ypt32 is incompatible with stable binding to the core. Transition to the open state of TRAPPII enables activated GTP-bound Rab11/Ypt32 to diffuse away from the active site chamber, and this TRAPPII monomer is available for another round of nucleotide exchange. This movie was made using Chimera. Note that morphing transitions are for illustrative purposes and do not necessarily represent the actual conformation transition pathways.

## Notes

### Competing Interest Statement

The authors have declared no competing interest.

